# Viscoelasticity driven deformation dynamics of the substrate affect mechanically induced Ca^2+^ signals

**DOI:** 10.1101/2025.11.07.687272

**Authors:** H Peussa, S Peltola, A Tervonen, S Lehtimäki, M Kauppila, R Bhati, C Fedele, H Tran, E Mäntylä, A Priimägi, S Nymark, TO Ihalainen

**Affiliations:** Tampere University

**Keywords:** Viscoelasticity, substrate deformation, focal adhesion, calcium, ER, actin, PIEZO1, polyacrylamide, DR1-glass, light-controllable

## Abstract

The viscoelastic properties of the substrate are known to affect many aspects of cellular behavior from migration to differentiation. Cells are also known to sense deformations in the extracellular matrix (ECM). However, these mechanotransduction events have mostly been studied separately, and the emphasis has been on purely elastic materials although most tissues in the body are viscoelastic. We wanted to understand how the viscosity and elasticity of the substrate affect cells’ mechanosensation, i.e., how cells integrate passive (substrate viscoelasticity) and active cues (dynamic substrate deformation) into calcium transients. We used a light-controllable Disperse Red 1-glass (DR1-glass) coating together with a thin polyacrylamide (PAA) hydrogel with controllable viscoelasticity. This allowed us to generate local deformations in the cell-substrate interface on substrates with different viscoelastic properties. Inspecting the resulting calcium transients in a Madin Darby canine kidney type II (MDCK II) monolayer with the genetically encoded calcium indicator jRCaMP1b revealed that cells responded to deformation differently depending on the viscoelasticity of the substrate. On stiff elastic gels cells exhibited all in all the largest calcium responses as well as increased sensitivity to the magnitude of deformation, with larger deformations leading to stronger calcium signals, whereas on soft elastic and soft viscoelastic gels the magnitudes of deformation had less effect on the degree of calcium signals. Indeed, immunostainings showed that cells formed the strongest focal adhesions (FA) on stiff gels, albeit differences were surprisingly small. Instead, computational modeling revealed that forces generated at FAs were strongly dependent on the viscoelasticity of the substrate, with increased elasticity and decreased viscosity leading to larger forces. Moreover, viscoelasticity affected the dynamics of the force generation. On stiff elastic gels force increased in fast steps, whereas on soft gels the buildup was gradual and was further slowed down by increased viscosity. Surprisingly, experiments with PIEZO1 KO cells showed that the calcium responses to the substrate deformation did not require PIEZO1 channels. Instead, depleting the ER with thapsigargin (TG), depolymerizing the actin cytoskeleton with cytochalasin D (cyto D) and latrunculin A (Lat A), and inhibiting actomyosin II with y-27632, showed that the calcium was mainly released from the ER in an actin, but not actomyosin, dependent mechanism. The data therefore suggests that the forces subjected to FAs could be directly transmitted to the ER via an actin mediated tension that results in the opening of ER residing calcium release channels. Our results illustrate that the viscoelasticity of the cell niche controls cell behavior in two ways. Firstly, it affects how cells build adhesions to the ECM and thus the accumulation of mechanosensitive proteins. Secondly, it affects the dynamics of the deformations and forces that are sensed by FAs. In line with our previous findings, we show that the mechanically induced calcium responses are dependent on dynamics, with fast mechanical cues resulting in larger calcium responses. Moreover, our research suggests that cells may possess distinct response mechanisms that are selectively activated depending on the mechanical properties of their niche, and the type and dynamics of the mechanical stimulation.

## 1 Introduction

It is well established that the mechanical properties of the cell niche guide the behavior of cells, affecting, for example, their migration and differentiation, as well as pathological processes such as cancer development (Chaudhuri et al., 2020; Mai et al., 2024). Traditionally, research on cell-ECM interaction has focused on the elasticity of the substrate, simply by comparing cellular phenomena on soft versus stiff substrates. However, most tissues in the body are not purely elastic, but instead viscoelastic, i.e., the material resists deformation in a rate dependent manner (Chaudhuri et al., 2020). Interestingly, it has been shown that soft viscoelastic materials can momentarily appear as stiff substrates to cells, as the lag time in material deformation allows traction forces to be formed (Chaudhuri et al., 2015). A viscoelastic substrate may therefore be interpreted as soft or stiff depending on how well the time scale of the cellular process matches with the lag in material deformation (Gong et al., 2018). This illustrates that not only the mechanical, but also the temporal aspects of material properties influence cellular behavior.

Cells adhere to their surrounding substrate via integrins that form a link between the cytoskeleton and the fibers of the extracellular matrix (ECM). In FAs, the most prominent force-generating cell-ECM junctions, integrins are clustered and accompanied by multiple mechanosensitive proteins such as talin and vinculin that link integrins to the actin cytoskeleton, signaling proteins such as focal adhesion kinase (FAK) that modulate FA dynamics, and numerous other docking and regulatory proteins (Martino et al., 2018). Cells are known to detect and to adjust to the stiffness of their surroundings through FAs by increasing the traction forces in these junctions to match the resistance from the substrate (Discher et al., 2005; Elosegui-Artola et al., 2018). On stiff substrates, the increased tension between integrins and the ECM promotes the recruitment of additional FA proteins and actin into the junction. Actin is remodeled into thick stress fibers where interaction with myosin II enables traction forces to be generated. This acts as a positive feedback loop further increasing the FA assembly until a balance in force is reached. On the contrary, on soft substrates, the ECM complies with the traction forces generated by the cell, thus failing to activate FA strengthening, leading to disassembly of the junctions, less actin recruitment, and less traction force to be generated. Moreover, strengthening of FAs affects the activity of signaling molecules, thus leading to wider functional alterations in cell behavior. For example, on stiff substrates FAK has been shown to activate the Erk/MAPK pathway, which controls cell cycle and differentiation (Tumenbayar et al., 2025). Consequently, mechanical information from the substrate controls many fundamental cell processes (Iskratsch et al., 2014).

In addition to feeling the stiffness of the substrate, cells also sense dynamic cues such as stretch, compression, and shear. These dynamic cues can trigger acute responses, often via ion channels that can generate responses in the scale of milliseconds (Cox et al., 2019). In non-excitable cells such as epithelial cells, calcium ions have an important role as universal signaling molecules, controlling a myriad of cell processes including proliferation, apoptosis, and gene expression (Berridge et al., 2000). Moreover, calcium is required for the assembly and disassembly of FAs and for actomyosin contraction making it crucial in FA strengthening (Jetta et al., 2023; Yao et al., 2022). Indeed, a calcium signal is often the first response to mechanical stimulation. It can be triggered by various mechanosensitive (MS) ion channels found on the plasma membrane (PM) that allow an influx of extracellular calcium (Ranade et al., 2015). In MDCK II cells, for example PIEZO1 and some members of the Transient receptor potential (TRP) channel family including TRP vanilloid 4 (TRPV4), and TRP polycystin 2 (TRPP2, also known as PKD2) can generate a calcium influx in response to various types of mechanical stimulation (Coste et al., 2010; Rocio Servin-Vences et al., 2017; Song et al., 2024). Emerging data suggests that integrin mediated mechanical stimulation of the PM can also trigger a calcium release directly from the ER even without an initial extracellular influx of calcium (Kim et al., 2015). This suggests that mechanosensitive structures exist also at the ER. For example, the before mentioned TRPP2 has been shown to localize on the ER, and to be involved in mechanically induced calcium release from the ER (Song et al., 2024). In addition, recently, the inositol 1,4,5-trisphosphate receptors (IP3R), the main calcium release channels on the ER, as well as stromal interaction molecule (STIM) and Orai, the main components of the store operated calcium entry (SOCE) pathway, have been shown to require actin to function, thus raising the possibility for mechanosensitive regulation (de Souza et al., 2021; Thillaiappan et al., 2021). Moreover, STIM has been identified as an important ER-PM tether that is required for membrane tension to be transmitted to the ER (Chen et al., 2025). This demonstrates that cells are equipped with multiple layers of mechanosensitive structures that enable them to fine tune their response to different external mechanical cues.

The mechanisms that enable cells to sense substrate stiffness and substrate deformation are well known. However, these phenomena are usually studied separately, and, until lately, the focus has been on the elastic properties of the substrate with less focus on the viscosity. We therefore wanted to understand the combined effect of how cells respond to dynamic substrate deformations in different viscoelastic environments. To this end, we developed a system where a hydrogel layer can be deformed in the presence of living cells, using light as the actuator. This enables mechanical stimulation of living cells on substrates with different viscoelastic properties while imaging them simultaneously. To detect fast cellular responses, we used calcium imaging. Here, we show that the system enables robust deformations of the cell substrate in a micrometer scale, and that the viscoelastic properties of the substrate affect the degree of generated calcium dynamics in the stimulated cells. Differences in cell responses are explained by a combined effect of substrate deformation dynamics and strength of FAs, both of which were found to depend on the viscoelastic properties of the substrate.

## 2 Results

### 2.1 Construction of glass-DR1-PAA-fibronectin structures

It is well established that various cell types can sense the mechanical stiffness and viscous properties of the tissue and the surrounding ECM (Janmey et al., 2020) (Chaudhuri et al., 2020). Furthermore, cells are able to respond to nanoscale changes in ECM topography, indicating that they possess a highly responsive mechanosensing machinery (Peussa et al., 2023; Poole et al., 2014). We wanted to understand how the cells integrate multiple physical cues, and more specifically, how substrate stiffness and viscosity influence the sensing of small ECM deformations.

The experimental setup we developed here consisted of a polyacrylamide (PAA) hydrogel polymerized on a 250 nm thick Disperse Red 1 glass (DR1-glass) coating on a glass coverslip. DR1- glass, an azobenzene based molecular glass, undergoes cis-trans-conformational changes upon light stimulation (Kirby et al., 2014). These changes transiently liquefy the DR1-glass, leading to a lateral flow away from the illuminated region and when scanned with a laser beam, cause a controlled lateral displacement and piling of the material. Fluorescent beads (⌀ 0.2 µm) were embedded into the gel to visualize and quantify the subsequent gel deformation and thickness. These beads have negligible influence on mechanical properties of the gels and have been widely used e.g., in traction force microscopy (Plotnikov et al., 2014). The elastic modulus (E’) of the PAA gel was adjusted by varying acrylamide and bis-acrylamide ratios (Tse and Engler, 2010), whereas viscous modulus (E’’) was tuned by adding long linear PAA chains to increase internal friction of the polymer network (Charrier et al., 2020). The gel surface was coated with ECM protein fibronectin to promote adhesion of MDCK II epithelial cells (Figure 1A).

**Figure 1:**
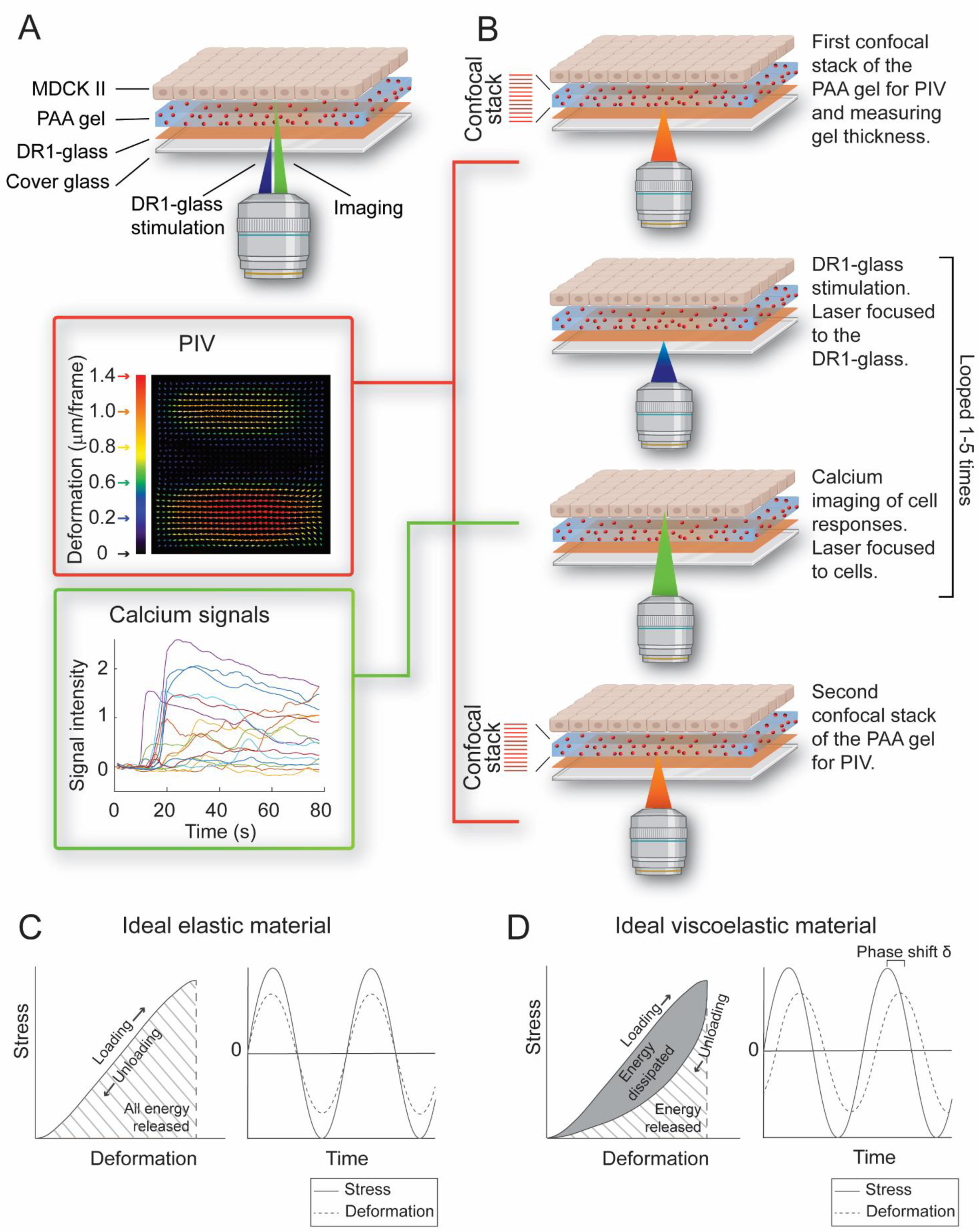
Schematics of the experimental setup. A) The setup consisted of a light controllable DR1-glass coating, a PAA gel layer with tunable elastic and viscose properties and MDCK II cells. Fluorescent beads withing the hydrogel layer enabled tracing gel deformation and measuring gel thickness. B) Experiments were carried out with a laser scanning confocal microscope. DR1-glass was stimulated by focusing the laser beam (488 nm) on the DR1-glass layer. Generated calcium dynamics in the epithelial monolayer were tracked with jRCaMP1b calcium indicator by focusing the 561 nm laser beam to the cell cytoplasm. Confocal stacks were taken of the fluorescent beads (647 nm) before and after the experiment. Gel deformation was determined based on bead displacement using particle image velocimetry (PIV) analysis. C) The stress-deformation curve and temporal relation between stress and deformation in an ideally elastic material. Deformation occurs simultaneously with stress without energy loss. D) The stress-deformation curve and temporal relation between stress and deformation in an ideally viscoelastic material. Part of the energy is stored within the structure of the material. This leads to a delay in the relaxation called the phase shift.

Experiments were performed with a laser scanning confocal microscope (LSCM) by alternating the laser wavelengths (647 nm for fluorescent beads in the PAA gel, 488 nm for DR1-glass stimulation and 561 nm for the calcium indicator jRCaMP1b) and by switching between the focal planes of the DR1-glass, PAA gel, and cells (Figure 1B). This configuration allowed us to quantify gel deformation and thickness from the confocal stacks, stimulate the DR1-glass layer, and record the calcium dynamics in the cells. The use of MDCK cells expressing genetically encoded Calcium indicator jRCaMP1b allowed the direct imaging of cytosolic calcium dynamics in response to substrate deformation.

In this study we used a soft elastic, a soft viscoelastic and a stiff elastic gel. In purely elastic materials, stress and deformation occur simultaneously (Figure 1C), whereas in viscoelastic materials part of the mechanical energy dissipated as internal friction, producing a time delay or phase shift between the applied stress and resulting deformation (Figure 1D) (Wilson, 2018). The elastic (E’) and viscous moduli (E’’) of the PAA gels were tuned as described above, allowing the construction of substrates with distinct mechanical properties and responses to physical stimulation.

### 2.2 Casting of thin PAA gels

We assumed that gel thickness strongly affects the extent of gel deformation, with thinner gels showing larger deformations on the top of the gel. However, too thin gels can allow cells to sense the rigid glass coverslip below the gel, affecting their physiology (Buxboim et al., 2010). Therefore, we optimized the gel casting procedure to achieve thin gels, aiming for 10 µm, with minimal variation. We performed this optimization using stiff elastic gels as a model. We placed weight on the uncoated top coverslip during polymerization to spread the gel evenly, and ⌀ 5 µm spacer beads were included to prevent over compression and ensure the minimal thickness of 5 µm (Buxboim et al., 2010). Because PAA gels can swell in aqueous conditions, the theoretical casting volume for 10 µm thick gels was adjusted empirically. During polymerization, the fluorescent beads accumulated near the top and bottom surfaces of the gels, enabling quantification of the gel thickness as a distance between the intensity peaks (Figure 2B).

**Figure 2:**
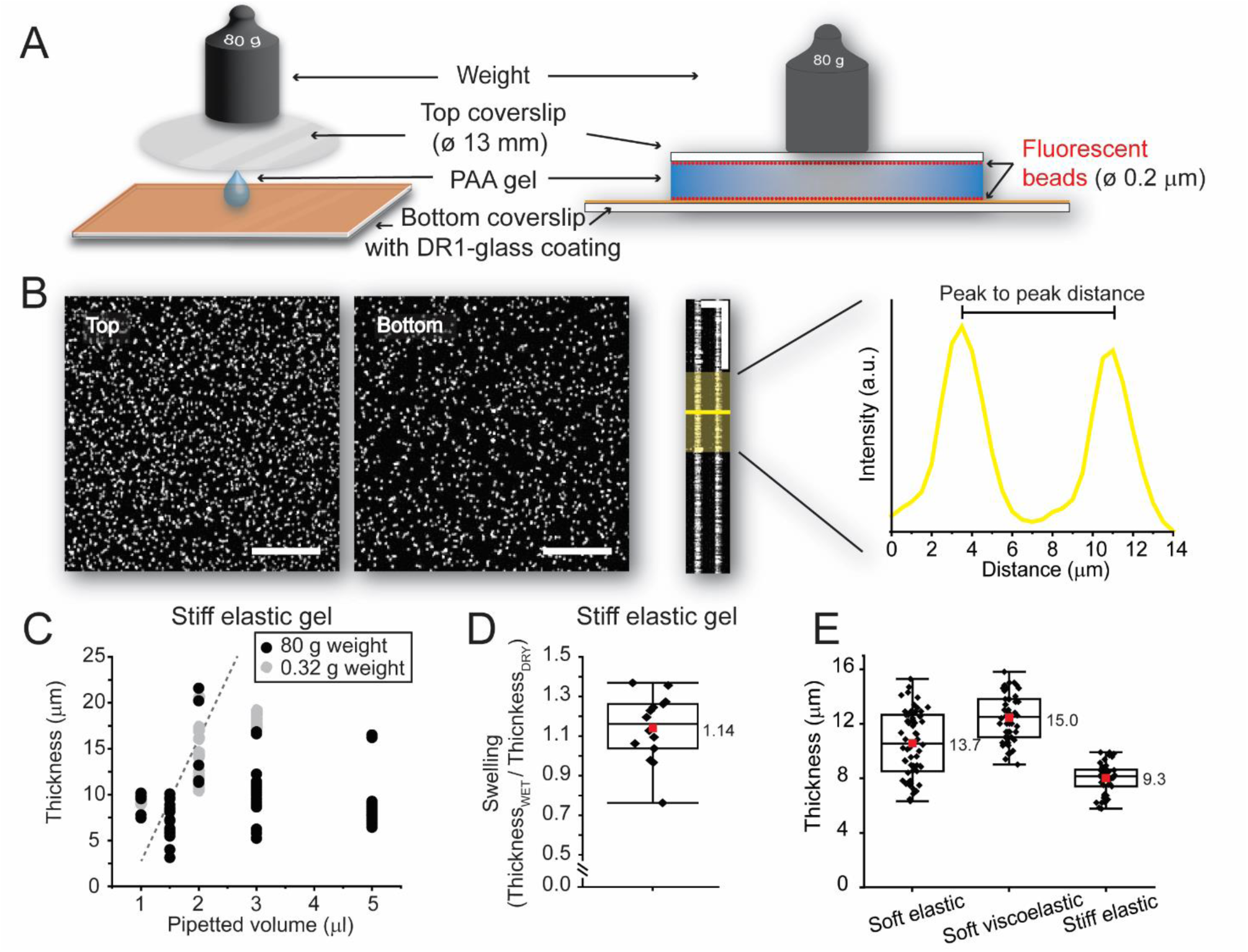
Gel thickness. A) Schematic of the gel casting procedure. Gel mixture containing fluorescent beads was pipetted onto the DR1-glasscoated coverslip. A passivated ⌀ 13 mm coverslip was placed on top and compressed with an 80 g weight for the duration of the polymerization (15 min). The fluorescent beads migrate toward the top and bottom of the gel layer. After polymerization the weight and top coverslip were removed, and samples were stored in PBS. B) Maximum intensity projections of the bead layers at the top and bottom of the gel and a side projection of the gel. Scale bars 10 µm. Gel thickness was measured by determining the distance between peak intensity signals arising from the fluorescent beads. C) Thicknesses reached with different pipetting volumes. Samples compressed a 0.32 g weight (grey dots) display a correlation between pipetting volume and gel thickness. Such a correlation is not evident in samples compressed with an 80 g weight (black dots). Dashed line shows the theoretical relationship between volume and thickness. D)The swelling of the PAA gels was determined by measuring the gel thickness immediately after polymerization and after 24 h incubation in PBS. Data in C and D is gathered only from stiff elastic gels. E) Thickness distributions of the three different gel types casted with the optimized procedure (5 µl gel solution pipetted, 80 g weight in top). In box plots, the whiskers determine the 5th and 95th percentiles, and the box determines the 25th and 75th percentiles. Means are marked with red squares, with mean values noted next to the bars. The vertical line represents the median value.

The pipetting volumes tested were 1 µl, 1.5 µl, and 3 µl, representing 7.5 µm, 11.3 µm, and 22.6 µm theoretical gel heights. When a light (0.32 g) weight was used on top of the coverslip, gel thickness scaled linearly with pipetted volume, whereas under heavier (80 g) weight, thickness became independent of the gel volume, even with 5 µl pipetted volume (Figure 2C). We also analyzed the swelling by imaging the gels immediately after polymerization and again after 24 h incubation in PBS. Thickness increased from 0 % to 40 % with no correlation to the pipetted volume (Figure 2D), suggesting local variability but minimal overall swelling in these conditions. Because pipetting small volumes of the viscous gel solution was technically challenging, we adopted the heavier 80 g weight, which allowed slightly excess of the gel mixture to spread evenly. Thus, here after, the gels were made by pipetting 5 µl of the gel solution that contained ⌀ 5 µm spacer beads and ⌀ 0.2 µm fluorescent beads, with 80 g weight applied during polymerization. Using this optimized method, the resulting thicknesses (Figure 2E) were 5.8 – 15.3 µm for the soft elastic gels, 9.0-16 µm for soft viscoelastic gels, and 5.8-9.9 µm for stiff elastic gels (n = 89, 67, and 69 respectively). Since gel thickness was quantified for each sample, the values could later be incorporated into gel deformation and Ca^2+^ response analyses to account the potential thickness-related effects.

### 2.3 Elastic and viscose properties of the PAA gels

To verify the mechanical properties of the gels we used, we determined their elastic and viscose moduli. This was essential, as in the publications where the recipes for the gels originate, the soft gels were characterized with rheological measurements (Charrier et al., 2020) whereas the stiff elastic gel with an atomic force microscope (AFM) (Tse and Engler, 2010). As AFM applies an indentation (Figure 3A) and rheological frequency sweep measurements torsion (Figure 3B), the two techniques measure different types of elastic moduli, making the values incomparable. Therefore, it was essential to analyze all gel types to get comparable values.

**Figure 3:**
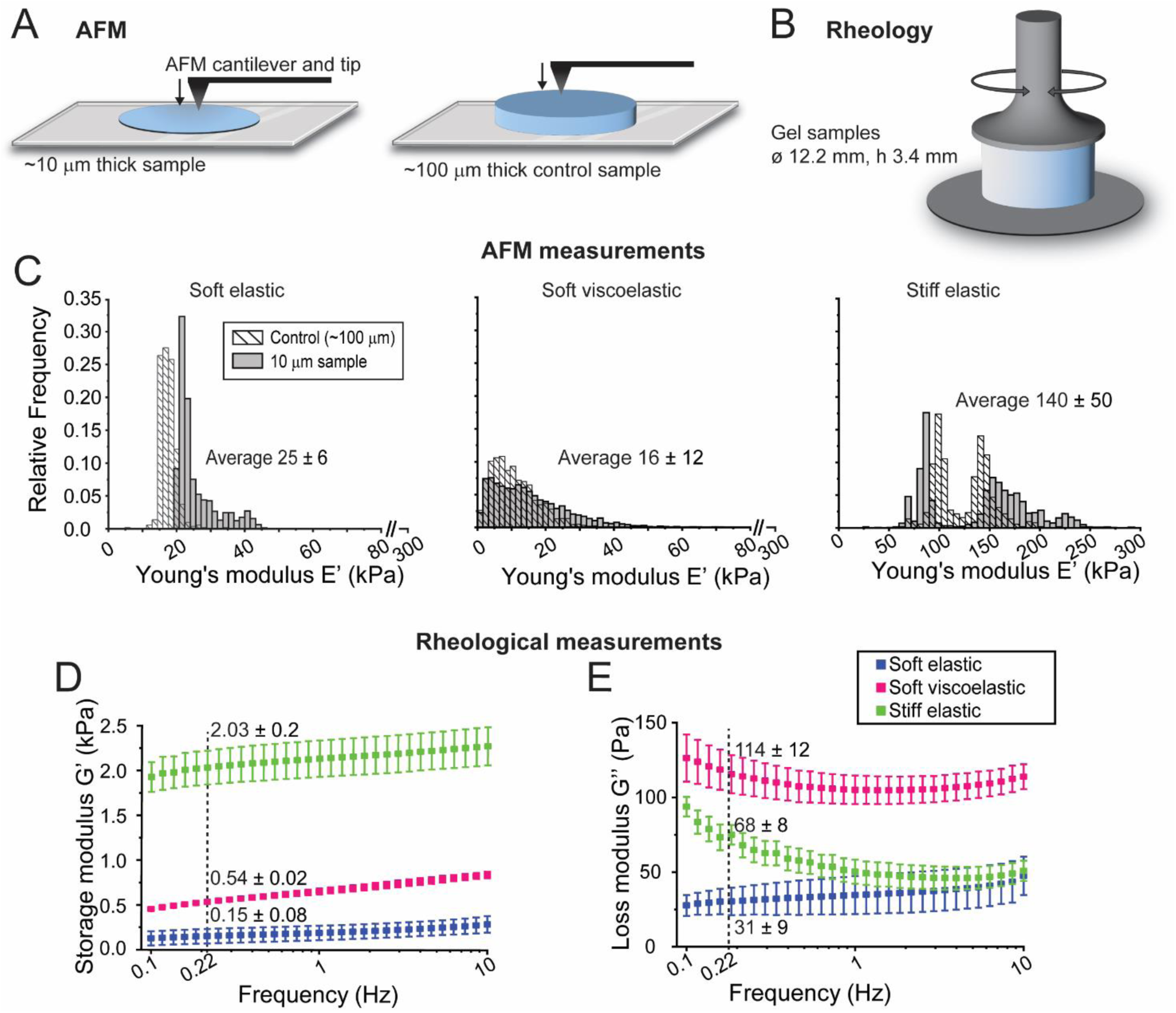
Characterization of the elastic and viscose moduli of the PAA gels. A) Schematic of the samples used for AFM measurements. Samples were manufactured as described in Figure 2 leading to ∼10 µm gel thickness. Similar samples were used in calcium imaging experiments, thus representing the mechanical environment sensed by cells. Thick control samples were prepared by pipetting 13 µl of gel solution and allowing gels to polymerize without any weight on top, resulting in ∼ 100 µm thick gels. B) Schematic of the rheological measurement procedure. The ⌀ 12.2 mm x h 3.4 mm gel samples were placed between plates and oscillated at an increasing frequency (0.1−10 Hz). C) AFM measurements were done on two occasions from triplicate samples. Controls were measured only on one occasion. Each sample was measured from multiple locations. Total number of measurements for soft elastic gels n = 4608, for soft viscoelastic gels n = 4352, for stiff elastic gels n = 4608 and for controls n = 512, n = 1536 and n = 512, respectively). Data was plotted into histograms of 20 bins using the range of 0− 150 kPa for soft gels and 0− 500 kPa for the stiff elastic gel. Thin samples are displayed as black and grey bars, and control data with the striped bars. The mean ± standard deviation of Young’s modulus values is displayed for each gel type. D) The mean ± standard deviation of values for storage modulus G’, and E) loss modulus (G’’). G’ describes the elasticity, and G’’ the viscosity, i.e., the internal friction. In D−E data from soft elastic gels shown in blue, soft viscoelastic in magenta and stiff elastic in green. Mean ± standard deviation of values is displayed for each gel type at the 0.22 Hz frequency data point, which matches the about 5 sec durations of each stimulation step.

We used AFM to determine local Young’s moduli (E’) in the PAA gels. These local measurements are especially interesting as they represent the mechanical environment sensed by the cells. Furthermore, as the sample is measured from multiple locations, the AFM measurements display the uniformity of elasticity within the gel samples. We measured multiple areas from 7 samples per gel type (3 + 4 independent technical replicates samples) using the force-volume mode. Additionally, we measured control samples with an approximate thickness of 100 µm. In these thick samples, the potential influence of the underlying rigid glass coverslip becomes negligible. The AFM measurements showed high inter- and intrasample variation (Figure 3C) indicating either uneven polymerization of the PAA gels, uneven mixing, or differences in sample thickness. The presence of the ⌀ 5 µm spacer beads may also have affected the measurements. However, we would expect similar high peaks in Young’s moduli in all sample types if in fact some of the measurement points coincided with the beads. We thus assume the effect of the spacer beads to be negligible. For soft elastic gels the control sample differed from the thin sample with slightly lower E’ values, whereas for the soft viscoelastic and stiff elastic gels, measurements from the thin and control samples fell within the same range (Figure 3C). This difference might fall within variation between samples, or it may suggest that the underlying glass coverslip does affect the mechanical status of the material. However, as the Young’s modulus of glass is in the GPa range and the measurements from the thin gels in the kPa range, the effect is minimal, indicating that the mechanical properties of the cell environment are governed by the gels.

Despite the variation, the results show that the soft gels depict similar ranges of E’, with mean values of 25 ± 6 kPa for soft elastic and 16 ± 12 kPa for soft viscoelastic gels, whereas for the stiff gel the mean Young’s modulus was 140 ± 50 kPa. This demonstrates that the soft gels have similar elasticities and that the stiff gel has on average a 7 times higher E’.

As traditional AFM cannot be used to measure viscose properties, we turned to rheology. However, rheological measurements can only be done from bulk material, so instead of measuring the ∼10 µm thick gels, we used gel samples with dimensions of Ø 12.2 mm x 3.4 mm. We did frequency sweeps ranging from 0.1 to 10 Hz with 1 % strain to determine the storage modulus (G’) and the loss modulus (G’’) of each gel type. The frequency sweep is the standard way to perform rheological measurements as viscosity is a time dependent modulus, making it important to measure the material’s behavior at different frequencies of stress. The stiff elastic gel had the highest G’ values throughout the frequency range, while the soft elastic and soft viscoelastic gel displayed lower G’ values. To compare individual values between the samples, we decided to focus on the 0.22 Hz datapoint as it matches the ∼5 sec duration of the DR1-glass stimulation loop. Moreover, similar time scales (5−10 sec range) have been reported for the dynamics of traction forces in FAs (Chan and Odde, 2008; Plotnikov et al., 2012), making this a relevant value to compare. Similarly to the AFM measurements, the soft elastic and soft viscoelastic gels had G’ values in the same range (0.15 and 0.54 kPa, respectively) while G’ for the stiff gel was on average 7-fold higher (2.03 kPa).

As expected, the loss moduli G’’ that tells of the viscosity of the material, was the highest for the soft viscoelastic gel, with a mean value of 114 Pa at 0.22 Hz (Figure 3E). At this frequency, the G’’ for soft elastic gel was 31 Pa and for stiff elastic gel 68 Pa.

Based on our characterization, the three different gel types had distinct viscoelastic properties that enabled us to analyze how mechanosensitivity of cells is affected by both the elastic and the viscose moduli of the substrate. The soft gels have similar elastic moduli but differ for their viscosity, allowing us to analyze the effects of the viscose modulus. Vice versa, the soft elastic and stiff elastic gels both have minimal viscose components, but vastly different elasticities, allowing us to compare the effects of the elastic modulus.

### 2.4 Magnitude and dynamics of gel deformation

Next, we wanted to characterize how stimulation parameters of the DR1-glass layer translate to gel deformation. As shown previously (Peussa et al., 2023), DR1-glass deformation scales with the 488 nm laser power and the number stimulation loops. To test the this behavior in our hydrogel system, we tested laser powers of 0.91 mW, 1.52 mW, 1.75 mW, and 2.59 mW, and stimulation loops of 1x, 5x and 5x5. Control experiments were performed using 488nm laser with 0 mW power, and 561 nm and 647 nm imaging lasers with 0.23 mW and 0.33 mW powers, respectively (maximum laser powers used in imaging) to confirm that the gel deformation originated from the DR1-glass activation. To rule out that the movement of the gel is generated by the DR1-glass movement instead of optical trapping of the fluorescent beads, we used flipped samples where the DR1-glass and the gel are on the opposing sides of the coverslip. By incorporating the DR1-glass layer beneath the sample we achieved a similar absorption spectrum as in normal samples, but as the gel was not in contact with the DR1-glass layer, no movement could transfer from the DR1-glass to the gel.

Gel deformation was determined by taking a confocal stack of the entire gel thickness before and after DR1-glass stimulation (Figure 4A) and comparing the bead positions using particle image velocimetry (PIV) (Tseng et al., 2012). By taking subsections of the confocal stack, we could separately analyze deformation at the bottom of the gel (directly attached to the DR1-glass layer) and at the top surface (where cells adhere). Increasing laser power and the number of stimulation loops increased deformation on both interfaces (Figure 4A−B). The largest deformations occurred when the 488 nm laser power was at least 1.52 mW with minimum 5 stimulation loops. No movement was observed in flipped sample controls, confirming that the gel displacement originated from the movement of the DR1-glass. Control imaging with 561 nm and 657 nm lasers produced only minimal movement confirming that the deformation was induced by 488 nm laser. We also performed control measurements to verify that the gel deformation remained stable beyond the observation window (confocal stacks taken before and after stimulation). Repeated PIV measurements after 5 min delay showed no change in the bead displacement, indicating that the gel neither continued to deform nor relaxed in this timeframe (Supplementary Figure 1A).

**Figure 4:**
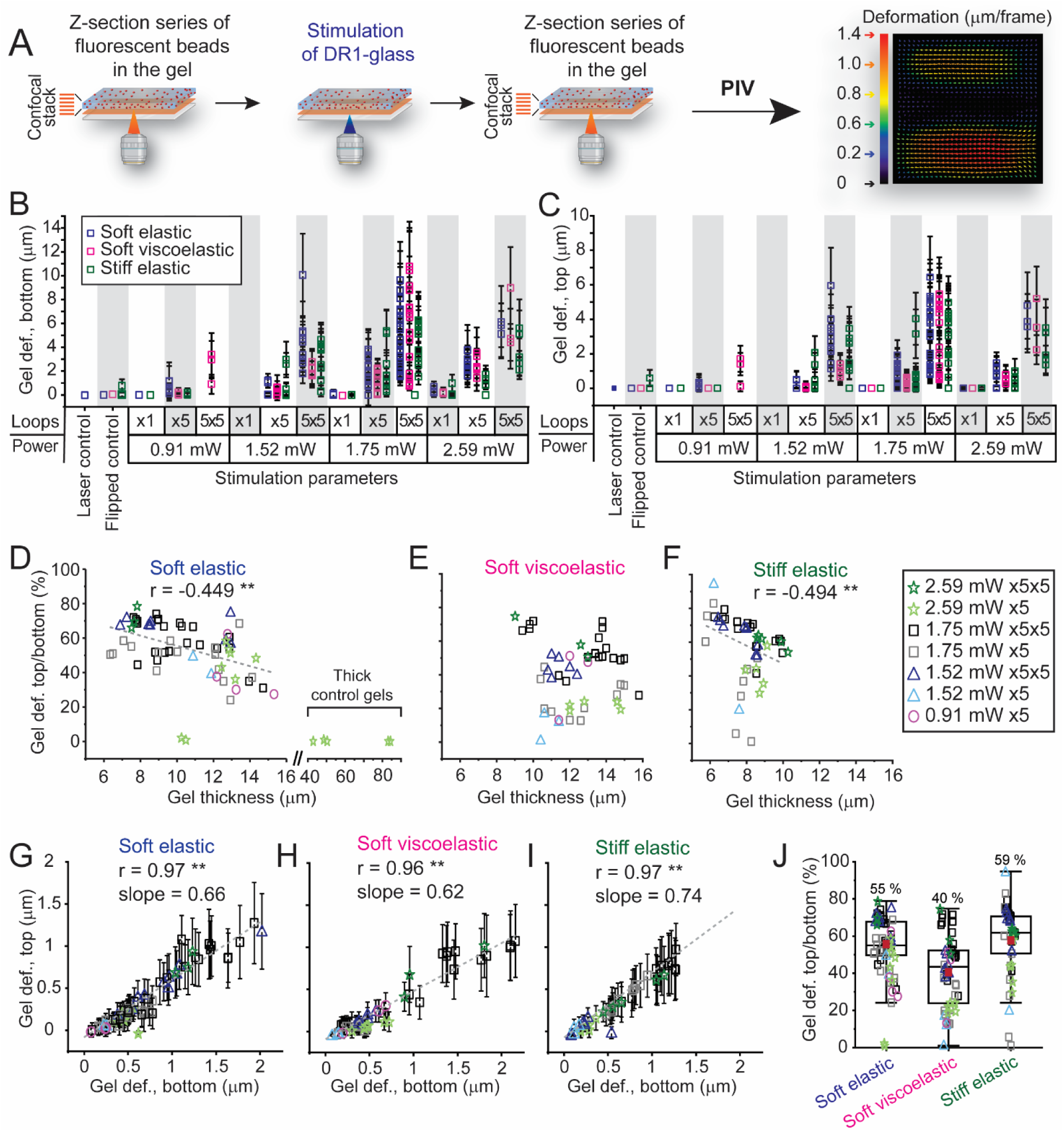
Gel deformation. A) Schematic of gel deformation measurements. Confocal stacks were taken of the gels before and after DR1-glass stimulation. The displacement of the fluorescent beads within the gel were analyzed with PIV. B) Mean ± standard deviation of gel deformation for soft elastic (blue), soft viscoelastic (magenta) and stiff elastic (green) gel at the bottom of the gel and C) top of the gel with different stimulation parameters (laser power and stimulation loops). In laser control, the DR1-glass was stimulated with the wavelengths and intensities used for imaging confocal stacks and calcium signaling (0.23 mW 561 nm laser and 0.33 mW 647 nm laser). In flipped control the gel was cast on the opposing side as the DR1-glass, resulting in identical absorption parameters while excluding the possibility of DR1-glass generated gel deformation. D−F) Effect of gel thickness to how well deformation reaches the top of the gel (mean deformation at top of the gel / mean deformation at bottom of the gel) in the different gel types. For the soft elastic gel, also thick control gels were included. Pearson’s correlation values are marked above plots. G-I) Mean ± standard deviation values at the top of the gel as a function of the mean deformation at the bottom of the gel. Pearson’s correlation values and sloped of linear correlation marked above plots. J) Percentage of preserved gel deformation (deformation at top of the gel / deformation at bottom of the gel). In D−M stimulation parameters are indicated with color and symbols (2.59 mW x5x5 = dark green star, 2.59 mW x5 = light green star, 1.75 mW x5x = black square, 1.75 mW x5 = grey square, 1.52 mW x5x5 = dark blue triangle, 1.52 mW x5 = light blue triangle, 0.91 mW x5 = magenta circle). In box plots, the whiskers determine the 5th and 95th percentiles, and the box determines the 25th and 75th percentiles. Means are marked with red squares, with mean values noted next to the bars. The vertical line represents the median value. *p < 0.05, **p < 0.01, and ***p < 0.001 by Student’s t test (D, F, G, H and I).

The variation in mean deformation at the bottom of the gel (i.e., directly on top of the DR1-glass layer) with similar stimulation parameters was large (Figure 4B). Consequently, also mean deformation on the top of the gel varied a lot (Figure 4C). We assumed that the thicker the gel, the less deformation would reach the top of the gel. Indeed, such correlation was observed in soft elastic gels (Pearson correlation r = -0.449**) and in stiff elastic gels (r = -0.494**) (Figure 4D, F). However, no significant correlation was detected for the soft viscoelastic gels (Figure 4E). In thick control gels that were over three times thicker, no deformation propagated to the top surface (Figure 4D), indicating that thickness influences the resulting deformation at the gel surface. However, within the thickness range used in our thin gels, thig effect remained minimal.

Moreover, the deformation at the top of the gel correlated strongly with the deformation at the bottom of the gel (Pearson correlation rsoft elastic = 0.775**, rsoft viscoelastic = 0.933**, rstiff elastic = 0.837**) (Figure 4G−I). On average, 54 % of the bottom deformation reached the top of the gel in soft elastic gels, 40 % in soft viscoelastic gels, and 58 % in stiff elastic gels (Figure 4J). Interestingly, in the soft gels the mean deformations at the bottom of the gel reached values up to approximately 2 µm, whereas in stiff gels the highest values were approximately 1.4 µm (Figure 4F−H). However, the percentage of deformation that reached the top of the gel was highest in the stiff gels (Figure 4J), suggesting that the stiff gel offers greater resistance to DR1-glass movement than soft gels, but transfers the deformation to the top surface more efficiently. This is seen also in the steeper slope of the regression line for the stiff elastic gels (Figure 4G−I).

Lastly, to understand how the material behaves under repeated stimulation, we performed PIV analyses on consecutive stimulations and followed the deformation across the loops, each consisting of five stimulation steps (Figure 5A). In the soft elastic gel, the deformation was greatest in the first stimulation loop and diminished in the following loops (Figure 5B), while in the soft viscoelastic and stiff elastic gels the deformations per loop were smaller but more consistent (Figure 5 C−D). In all gels, the deformation accumulated in a relatively linear fashion with some exceptions where the deformation saturated after three loops (Figure 5E−G). As the material behaves in a systematic manner from loop to loop, it suggested that cells will similarly experience consistent mechanical stimulation during repeated activations (Supplementary Figure 1B−C).

**Figure 5:**
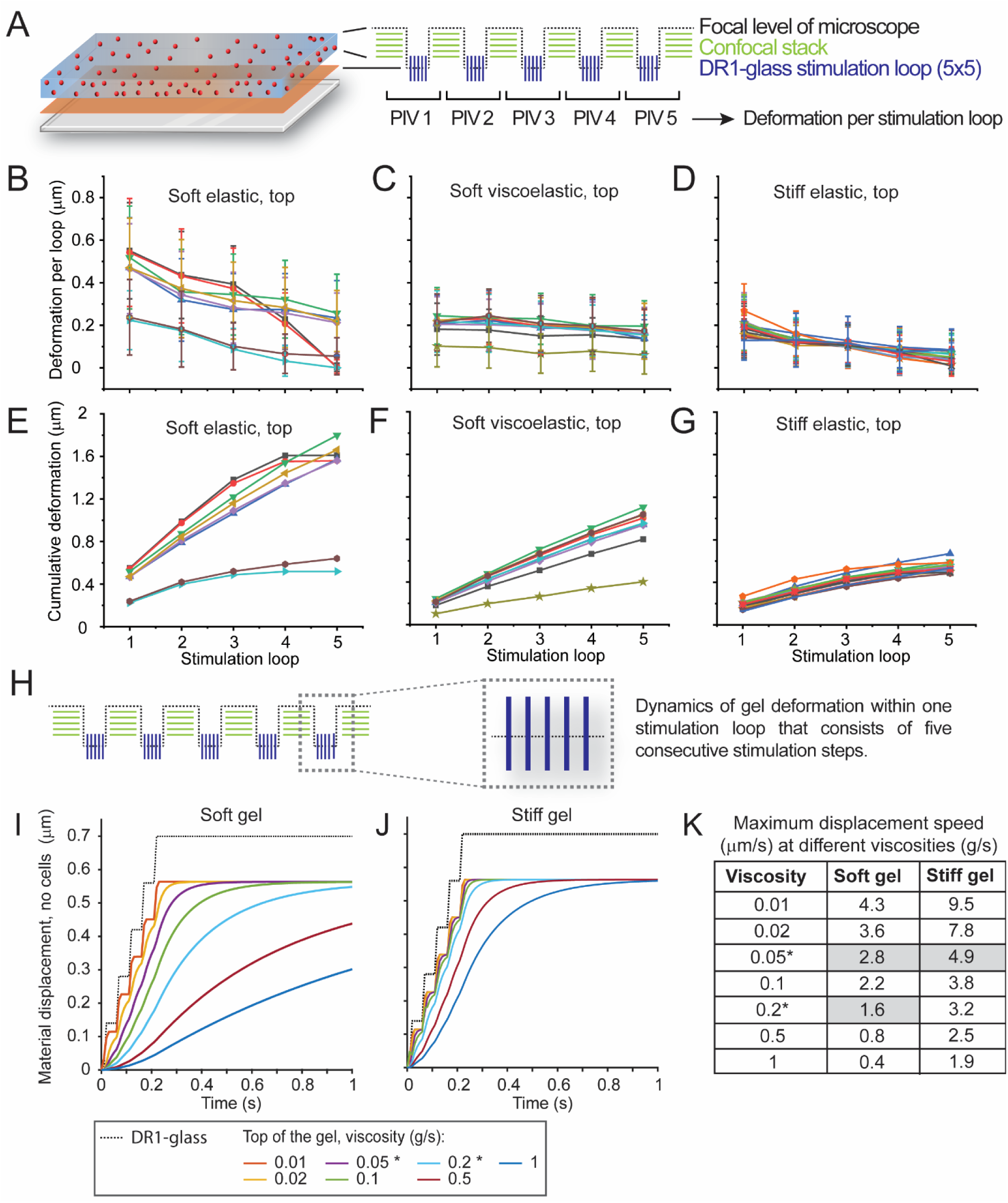
Cumulative deformation and dynamics of the gel deformation. A) Schematic of the experiment. A confocal stack of fluorescent beads within the gel layer was taken before stimulation and after each stimulation loop. Each loop consisted of 5 stimulation steps with 1.75 mW laser power. B−D) Mean ± standard deviation of deformation at the top of the gel after each stimulation loop for soft elastic, soft viscoelastic and stiff elastic gels. E−G) Cumulative deformation from loop to loop for each gel type. H) Gel dynamics within one stimulation loop (consisting of 5 consecutive stimulation steps) were analyzed by computational modeling. I) The movement of DR1-glass (black dashed line) and the surface of the soft gel and J) stiff gel with different viscosities (g/s). *For later simulations, 0.2 g/s was chosen to represent viscoelastic gels and 0.05 g/s elastic gels. K) Maximum displacement speeds in soft and stiff gels with different viscosity values.

While PIV measurements determine the absolute deformation of the gel, they only capture bead displacement before and after stimulation and do not provide information on the dynamic behavior of the material during the stimulation process. Because our system does not enable simultaneous DR1-glass stimulation and imaging of the substrate dynamics across multiple focal planes, and is further limited in temporal resolution, we turned to computational simulations to understand the immediate dynamics of the gel movement occurring during the stimulation loop. We used a simple gel model allowing adjustment of gel stiffness and viscoelasticity, as previously described by Tervonen et al. (2023).

As experimental data showed that gel deformation was relatively similar from stimulation loop to loop (Figure 5B−G), we concluded simulating the gel displacement during one loop (consisting of five repetitive stimulation steps, Figure 5H) was sufficient. Figures 5I and J show the gel displacement as a function of time, with the DR1-glass layer shown with a dotted line and the top of the gel with colored lines indicating different amounts of viscosity described by a so-called damping constant in the model. Since the observed deformation of even the viscoelastic gel occurs entirely in the time window between the stimulus and imaging (Supplementary Figure 1A), we chose a value of 0.2 g/s for the damping constant for the viscoelastic gel. To match the 4-fold difference detected in loss modulus in soft elastic and soft viscoelastic gels (Figure 3E), we approximated the damping constant of the elastic gels to be 0.05 g/s. The simulations show very different gel deformation dynamics depending on the viscosity, especially in the soft gels (Figure 5I−J, viscosity values 0.2 versus 0.05 g/s). We also calculated the maximum material displacement speeds and saw that in soft gels the maximum movement speed decreases from 2.8 µm/s to 1.6 µm/s as the viscosity increases from 0.05 to 0.2 g/s (Figure 5K). Meanwhile, the stiff elastic gels (viscosity 0.05 g/s) maximum displacement speed was 4.9 µm/s. The simulations show that deformation dynamics were affected by both elasticity and viscosity of the gel, with stiff elastic gels exhibiting the fastest and soft viscoelastic gels exhibiting the slowest dynamics.

### 2.5 Viscoelastic properties of the substrate affect cell-ECM adhesions

While it is known that cells form stronger focal adhesions (FAs) on stiff substrates (Yeh et al., 2017a; Zhou et al., 2017), the effect of substrate viscosity on cell-ECM junctions is less known. To address this, we performed immunostaining to compare the FAs and actin organization in cells grown on different gel types. For imaging FAs, we used antibodies for phosphorylated FAK (pFAK) and vinculin, and for imaging the actin cytoskeleton, we used phalloidin staining. Moreover, we have previously shown that mechanosensitive PIEZO1 channels are located on the basal side of cells cultured on hard substrates, where they act as key mediators of basal shear sensing (Peussa et al., 2023). Thus, we sought to determine whether the viscosity and elasticity of the substrate influence this localization by using a cell line expressing PIEZO1-HaloTag® fusion protein.

We compared the FAs in cells grown on different gel types based on pFAK and vinculin immunostainings (Figure 6A). To quantify the variation in FAs and evaluate their maturation across different gel types, we segmented the pFAK and vinculin signals. FA maturation leads to larger and more longitudinal FAs as well as increased clustering and recruitment of vinculin (Zimerman et al., 2004). We thus analyzed the area, elongation (determined as the eccentricity, which describes how the shape of the object deviates from a perfect circle, with 0 indicating perfect circle and 1 a line) and clustering (determined by measuring the distance to the first neighbor with shorter distance interpreted as increased clustering) of pFAK and vinculin. Based on both antibodies, the mean area of FAs was the smallest on soft elastic gels and largest on the stiff elastic gels (Figure 6B). Histograms also showed that on soft elastic and soft viscoelastic gels most FAs have small areas, whereas on stiff gels the distribution spreads also to larger areas (Supplementary Figure 2A-B). Due to the sample size, statistically significant differences were also found in the elongation of FAs, with most elongated structures found on soft elastic gels with both antibodies (Figure 6C), but the absolute differences were negligible. Lastly, both pFAK and vinculin showed the highest clustering on stiff elastic gels (Figure 6D). Visual inspection confirmed that large, elongated FAs were most common on stiff gels (Supplementary Figure 2).

**Figure 6:**
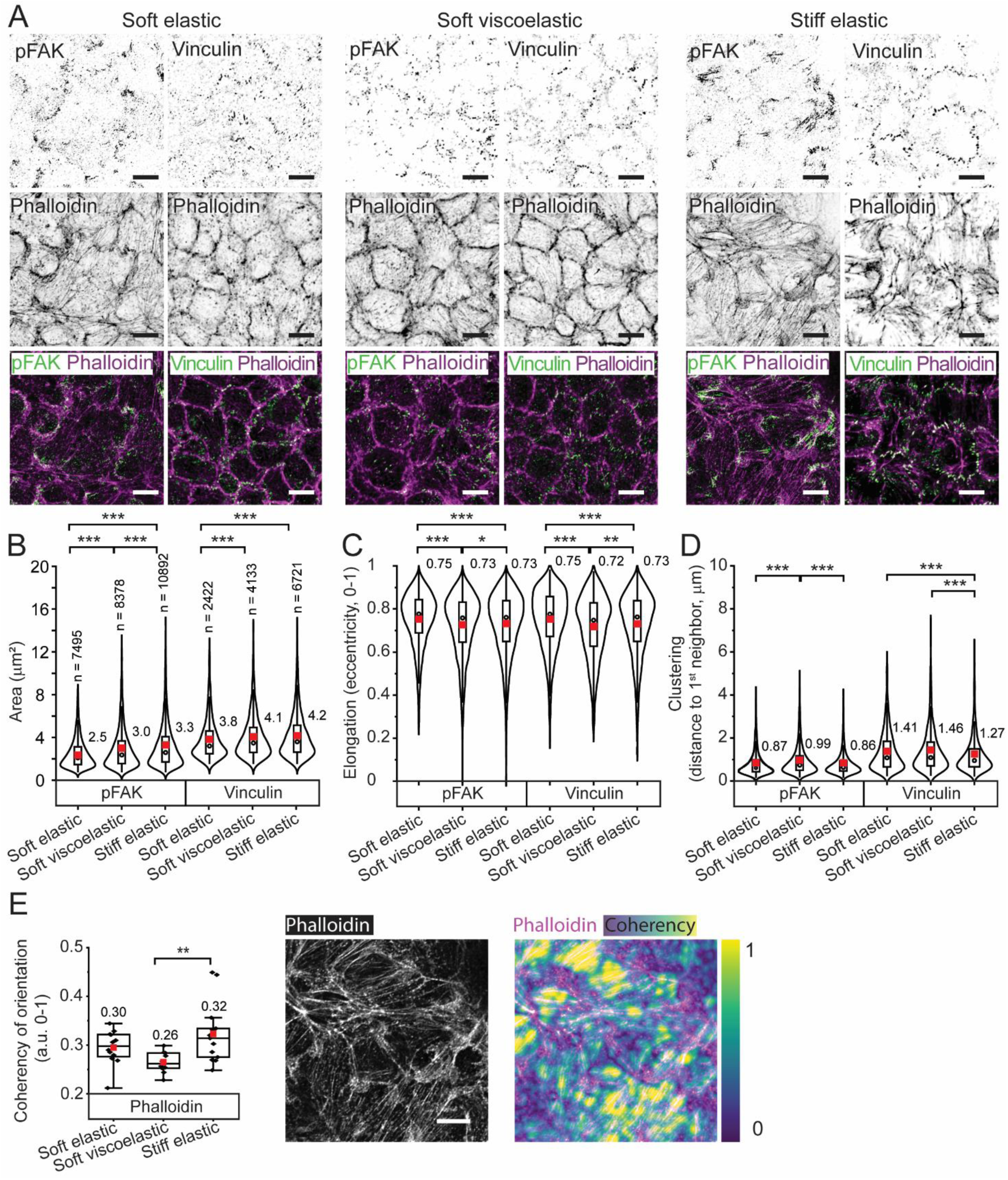
Focal adhesions and PIEZO1 channels. A) pFAK and vinculin stained together with phalloidin on soft elastic, soft viscoelastic and stiff elastic gels. Composites show pFAK and vinculin in green, phalloidin in magenta, and overlapping signal in white. Scale bars 10 µm. B) Analysis of the area (µm2), C) elongation (measured as eccentricity, a.u. between 0 and 1, where 1 signifies a line and 0 a circle), and D) clustering (measured as distance to first neighbor in µm) for segmented FAs based on pFAK and vinculin signal. E) Left, coherency of orientation (a.u. between 0 and 1, where 1 signifies perfect alignment and 0 isotropy) of phalloidin on soft elastic, soft viscoelastic and stiff elastic gels. Right, example image of phalloidin staining and a composite showing phalloidin (magenta) and the coherency (spectrum from violet to yellow). Scale bar 10 µm. In B-D, the violin displays the density plot of data points (Kernel distribution). In the inner box, the whiskers determine the 1.5x interquartile range, the box determines the 25th and 75th percentiles and the black circles show medians. Means are marked with red squares, with mean values noted next to the bars. In B the sample sizes are marked above the plots. *p < 0.05, **p < 0.01, and ***p < 0.001 by one-way ANOVA with Bonferroni Post Hoc test (B-D) or Kruskal-Wallis (E).

The quantification of the immunostainings supports what was detected with visual inspection: FAs tend to be more mature on stiff gels than on soft elastic and viscoelastic gels. Although the effect was more subtle than we had expected, our results align with existing literature showing how increased substrate stiffness enforces FA assembly (Yeh et al., 2017b; Zhou et al., 2017).

Actin cytoskeleton labeling with phalloidin revealed clear and more organized stress fibers in cells grown on stiff elastic gels than on soft gels, consistent with the previously reported higher cell-ECM adhesion forces on stiffer substrates (Doss et al., 2020). Since stress fibers often connect mature FAs, their emergence is a sign of FA assembly. We thus analyzed the coherence of orientation in the basal actin signal and saw that the stress fiber organization differed between soft viscoelastic and stiff elastic gels, with the highest mean coherency of orientation on stiff elastic gels (Figure 6E). This suggests that basal actin fibers are more aligned on stiff gels, which is in line with previous data, suggesting that cells formed stronger FAs on the stiff gels.

Lastly, we studied PIEZO1-channel distribution on the three tested gel types using a MDCK cell line expressing Halo-tagged PIEZO1 (Supplementary Figure 3). The confocal imaging showed that PIEZO1 channels were distributed on the basal side of cells and at cell vertexes on all three gel types. PIEZO1 has been proposed to participate in sensing substrate stiffness and to colocalize with FAs (Jetta et al., 2023; Yao et al., 2022). Therefore, we quantified the colocalization between PIEZO1 and the pFAK signal. However, both Pearson’s correlation coefficient (Figure 6G) and the Manders coefficient for PIEZO1 overlap with pFAK (Figure 6H) suggested that no colocalization occurs between PIEZO1 and pFAK in any of the gel types. Although total internal reflection fluorescence (TIRF) microscopy of purely basal PIEZO1 and pFAK in cells grown on glass substrates revealed subtle colocalization, this technique could not be used with gels (Supplementary Figure 3G). We also segmented the PIEZO1 images and quantified the total area of signal, area of individual clusters, the distance to the closest neighbor, and the number of clusters per image (Supplementary Figure 3C-F), but found no biologically meaningful differences between gel types. We therefore conclude that with the available imaging techniques, PIEZO1 expression and localization were similar on all gel types.

In conclusion, despite the significant variation, we detected that mature, strong FAs were more commonly found on stiff elastic gels. Moreover, on stiff gels the actin cytoskeleton formed stronger stress fibers indicating higher cortical tension, which is in line with increased FA strength. This agrees with the known phenomenon where increased substrate stiffness leads to stronger cell-ECM adhesions (Yeh et al., 2017b; Zhou et al., 2017). Lastly, we showed that PIEZO1 channels were found on the basal side of cells on all gel types, making them a plausible candidate for mechanosensation of substrate deformation.

### 2.6 Dynamics of deformation and FA strength affect mechanosensation

To compare how cells responded to substrate deformations induced by our setup, we followed cellular calcium activity, which is known to response rapidly to mechanical stimulation (Douguet and Honoré, 2019). We cultured MDCK cells stably expressing jRCaMP1b calcium indicator (Dana et al., 2016)) on the three different gel types. To capture baseline calcium levels, 10 frames were imaged before mechanical stimulation. Five stimulation loops were alternated with two frames of imaging to capture the gradual rise in intracellular calcium and finally a 60 frame timelapse of the cells was acquired to follow calcium activity after the stimulation (Figure 7A). The stimulation ROIs consisted of narrow lines, a large area and an unstimulated field in the middle to determine whether cell responses depended on the fraction of cell area exposed to stimulation. All cells in the field of view were nevertheless contacted by the stimulated ROI. The cells responded by raising their cytoplasmic calcium levels, although the magnitude and dynamics of the responses varied (Figure 7B). We didn’t observe any correlation between the stimulated cell surface area and cell responses, indicating that even a small stimulation area was sufficient to trigger a response (Supplementary Figure 4).

**Figure 7:**
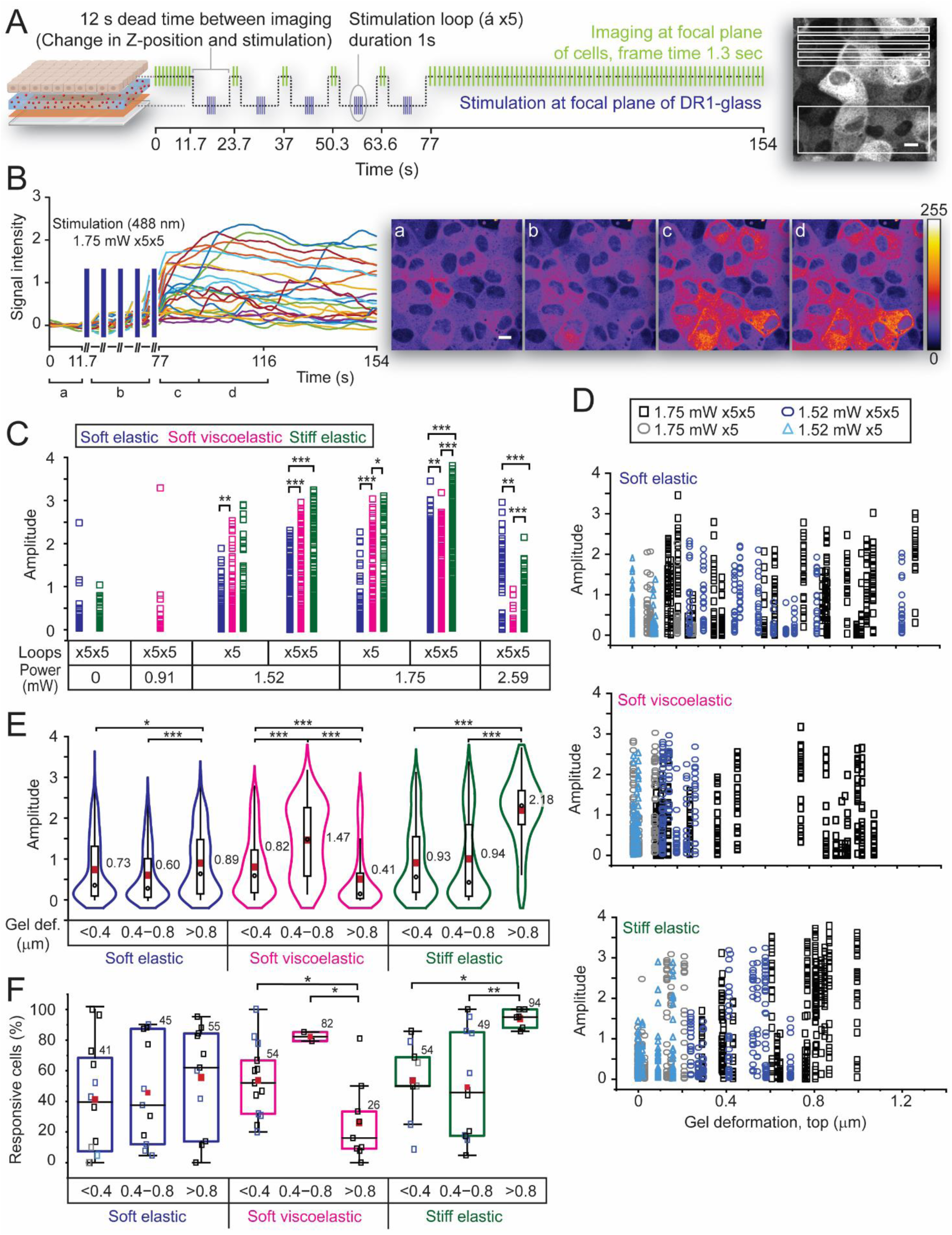
Calcium dynamics. A) A schematic representation of the imaging procedure (left). Ten frames of baseline activity were imaged before any mechanical stimulation. During imaging the laser beam (561 nm) was focused on the cell layer. For stimulation the focus was lowered to the DR1-glass layer, and 5 consecutive stimulations steps were performed. Focus was shifted back to the cell layer, and two frames were imaged. This was repeated 4 more times, followed by a longer (60 frames) time lapse at the end of the experiment. Imaging is indicated with light green lines and stimulation with blue lines. On the right, the stimulated ROIs are shown in white, scale bar 10 µm. B) Left, an example of the recorded calcium signals on a soft elastic gel, using 1.75 mW laser with x5x5 stimulations. Traces from 24 cells are normalized to zero with the baseline activity. Right, maximum intensity projections of the time-lapse data displaying the changes in calcium indicator intensity (black areas indicate low calcium concentration and yellow to white areas indicate high calcium concentration). The image data corresponds to the plotted data on the left and shows changes in jRCaMP1b signal intensity during (a) the ten baseline frames, (b) the stimulation steps, (c) the 10 frames directly after stimulation and (d) the following 20 frames. C) Calcium signal amplitudes generated with different stimulation parameters on the different gel types. D) Calcium signal amplitudes as a function of gel deformation. E) Violin plots showing calcium signal amplitudes at different gel deformation ranges (< 0.4 µm, 0.4−0.8 µm and > 0.8 µm) for each gel type. Box plots showing F) mean gel deformations that led to high amplitudes (amplitude > 2) in the different gel types, and G) percentage of responsive cells (amplitude > 0.5) at the different gel deformation ranges on the different gel types. In C, E, F and G, the different gel types are shown in blue (soft elastic), magenta (soft viscoelastic), and dark green (stiff elastic). In D, F and G the stimulation parameters are indicated with different colors (black square = 1.75 mW x5x5, grey ellipse = 1.75 mW x5, blue ellipse = 1.52 mW x5x5 and light blue triangle = 1.52 mW x5). In the violin plot, the violin displays the density plot of data points (Kernel distribution) and in the inner box, the whiskers determine the 1.5x interquartile range, the box determines the 25th and 75th percentiles and the black circles show medians. In box plots, the whiskers determine the 5th and 95th percentiles, the box determines the 25th and 75th percentiles, and the vertical lines show the medians. In all plots, means are marked with red squares, with mean values noted next to the bars. *p < 0.05, **p < 0.01, and ***p < 0.001 by one-way ANOVA with Bonferroni Post Hoc test (C, E) or Kruskal-Wallis (F).

Based on the gel deformation characterization, the 5x and 5x5 stimulations protocols (Figure 4) were used in the calcium-signaling experiments. Two main trends emerged. First, the amplitudes increased with the number of stimulation loops and with the increase in the stimulation laser power up to 1.75 mW (Figure 7C), indicating that the calcium response depends on the magnitude of the deformation (Figure 4C). Second, cells on stiff gels exhibited the largest calcium-signal amplitudes, suggesting that also FA maturity (Figure 6) and/or deformation dynamics (Figure 5I−K) influence cell responses.

To analyze how the deformation magnitude affected calcium responses, we plotted signal amplitudes against mean deformations. However, no significant correlation was detected between calcium signal amplitude and mean gel deformation (Figure 7D). To enable a more robust comparison between deformation and cell response, we classified the data into three categories based on the mean gel deformation: < 0.4 µm, 0.4−0.8 µm, and > 0.8 µm. In these categories, calcium signals amplitudes rose modestly with the increasing deformation in the soft elastic samples but increased substantially in the stiff samples (Figure 7E). However, on soft viscoelastic gels no clear trend between deformation and calcium response was detected (Figure 7E). The differences between calcium responses were the most striking with large gel deformations (> 0.8 µm): on the soft elastic gels the mean amplitude was 0.89 and on stiff elastic gels 2.18. This indicates that while larger deformations elicit stronger calcium responses, deformation magnitude alone does not explain the entire effect. The viscoelastic properties of the gel are an additional co-regulator of the response.

Since the calcium signal amplitudes showed large variance, we decided to analyze also the number of responsive cells per field of view (i.e. cells whose amplitude exceeds the levels of baseline activity, amplitude > 0.5). The data showed the same trend as the calcium signal amplitude, on soft elastic gels the percentage of activated cells increased modestly as the gel deformation increased, on stiff gels the increase was much larger, and on soft viscoelastic gels we did not detect any clear trend (Figure 7G). The effect was again most apparent with large deformations, as 55 % of cells on soft elastic gels and 94 % of cells on stiff gels were responsive.

Next, we wanted to understand the differences in the mechanical stimulation which could lead to the detected differences in the cellular responses. We used the computational model to simulate how forces emerged at the cell-gel interface. Based on our FA quantifications (Figure 6) and literature (Yeh et al., 2017b; Zhou et al., 2017), the constant describing FA substrate binding strength was assumed to be higher on the stiff gels than on the soft elastic and soft viscoelastic gels (Figure 8A). The simulated gel deformation lead to increased forces experienced by the FAs (Figure 8B). Next, we simulated the entire stimulation procedure (5 stimulation loops, each consisting of 5 stimulation steps) and plotted the maximum FA forces in the three different deformation rages (< 0.4 µm, 0.4−0.8 µm and > 0.8 µm) (Figure 8C). Interestingly, the simulations showed very small differences in maximal FA forces between soft elastic and stiff elastic gels even at large deformations. This suggested that the differences in calcium response are not explained by forces experienced by FA due to the gel deformation. Next, we simulated the gel deformation dynamics during a single stimulation loop by setting the deformation to 0.5 µm (Figure 8D). As expected, the plateaus of the FA forces were again similar on the stiff gel and soft elastic gel and notably smaller on the soft viscoelastic gel, but the dynamics of the force build-up between the gels were different. On stiff gels the force increased in step-like manner, whereas on the soft gels, forces increased smoothly throughout the stimulation steps (Figure 5I–K). Thus, the experimental data and simulations suggest that the mechanically induced calcium responses depend on the magnitude and speed of the material deformation, and the strength of FAs, all of which are directly regulated by the viscoelastic properties of the substrate.

**Figure 8:**
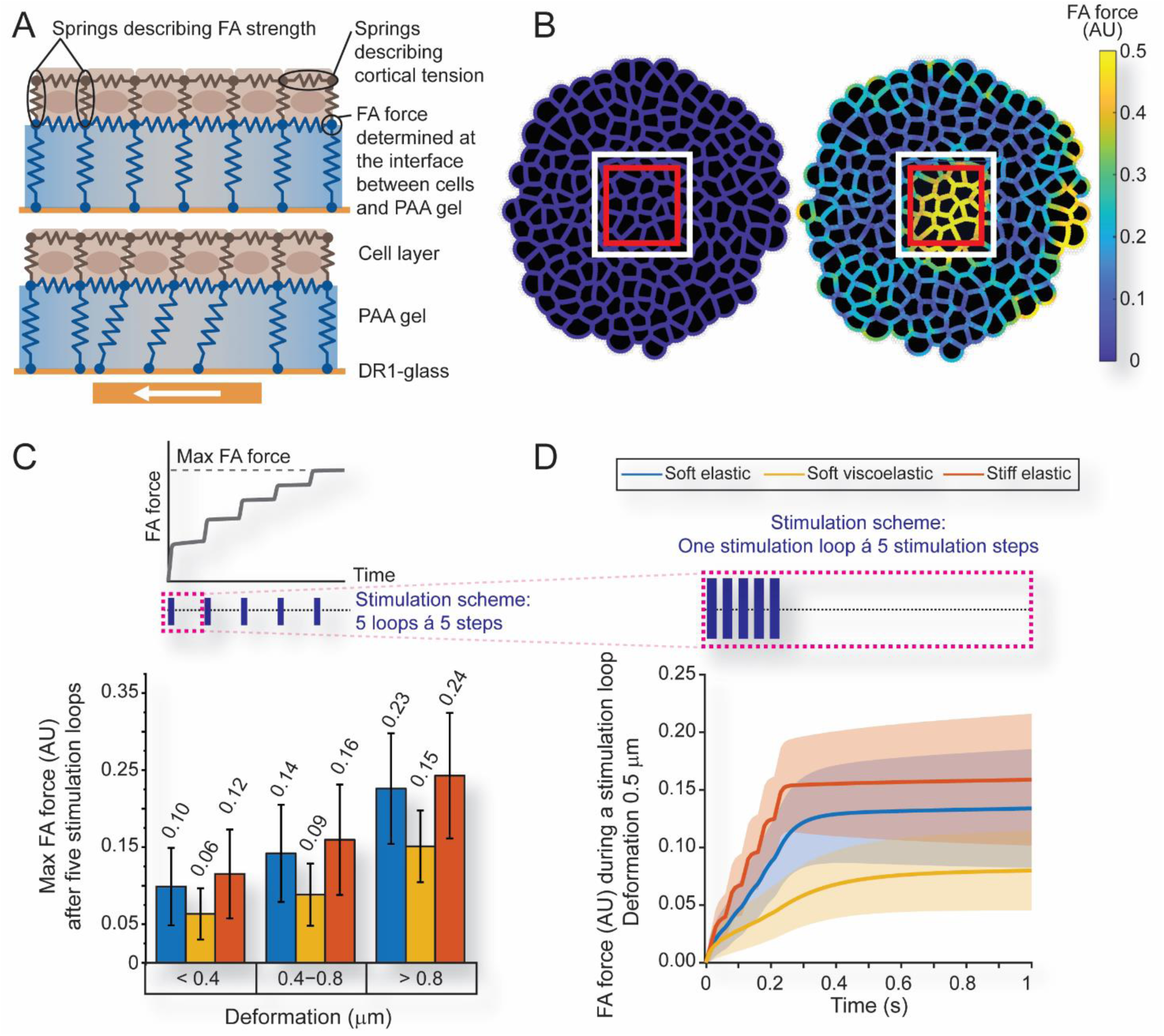
Simulations of force generation at the cell-ECM interface. A) Schematic of the model. The brown springs describe the FA strength (vertical) and cortical tension (horizontal) within the cells. The blue springs describe the elasticity of the PAA gel. The FA forces were calculated from the interface between the cells and the gel. B) Cell layer showing the borders of cells, the stimulated ROI (white square), and area where force measurements were done (red square). C) Above, schematic illustration describes the maximum FA force that was reached after 5 stimulation loops (stimulations marked with blue vertical lines, first loop is marked with a magenta dashed line). Below, mean and standard deviation of the max FA force on different gels within the different deformation ranges (< 0.4 µm, 0.4−0.8 µm and > 0.8 µm). D) Above, the stimulation scheme during one stimulation loop (magenta dashed line), consisting of five stimulation steps (shown as blue vertical lines, time corresponds to the plot below). Below, mean and standard deviation of the FA force as a function of time during one stimulation loop when gel deformation was set to 0.5 µm on all gel types. Forces were calculated from each cell inside the stimulated area. In C and D, soft elastic gel is shown in blue, soft viscoelastic in yellow, and stiff elastic in orange.

### 2.7 Calcium signals are released from the ER via actin, independent of PIEZO1 channels

Finally, we wanted to gain insight into the mechanism behind the detected calcium transients. Based on our previous findings (Peussa et al., 2023), the channels’ localization at the cell-ECM interface on all gel types (Supplementary Figure 2), and due to their known mechanosensitive nature (Coste et al., 2010), the PIEZO1 channel was our first candidate. Our hypothesis was that PIEZO1 channels near the FAs open in response to increasing FA forces, initiating a calcium influx that is subsequently amplified by calcium-induced calcium release from the ER. We therefore used MDCK PIEZO1 knockout (KO) cell line expressing the same jRCaMP1b calcium indicator and compared mechanically induced calcium responses to those of wild type cells. We used stiff elastic gels because their deformation dynamics and high FA strength generated the largest calcium responses. Surprisingly, we saw no decrease in calcium signal amplitudes in PIEZO1 KO cells, indicating that PIEZO1 channels did not play a significant role in detecting gel deformation (Figure 9A). As we had previously observed considerably reduced calcium signals in PIEZO1 KO cells in response to basal shear deformation, we confirmed these results (Supplementary Figure 5C). We also again verified the PIEZO1 knockout by adding Yoda1 to the cells and observed only a negligible small rise in calcium compared to WT cells, thus confirming the lack of PIEZO1 functionality in our KO cell line (Supplementary Figure 5D).

**Figure 9:**
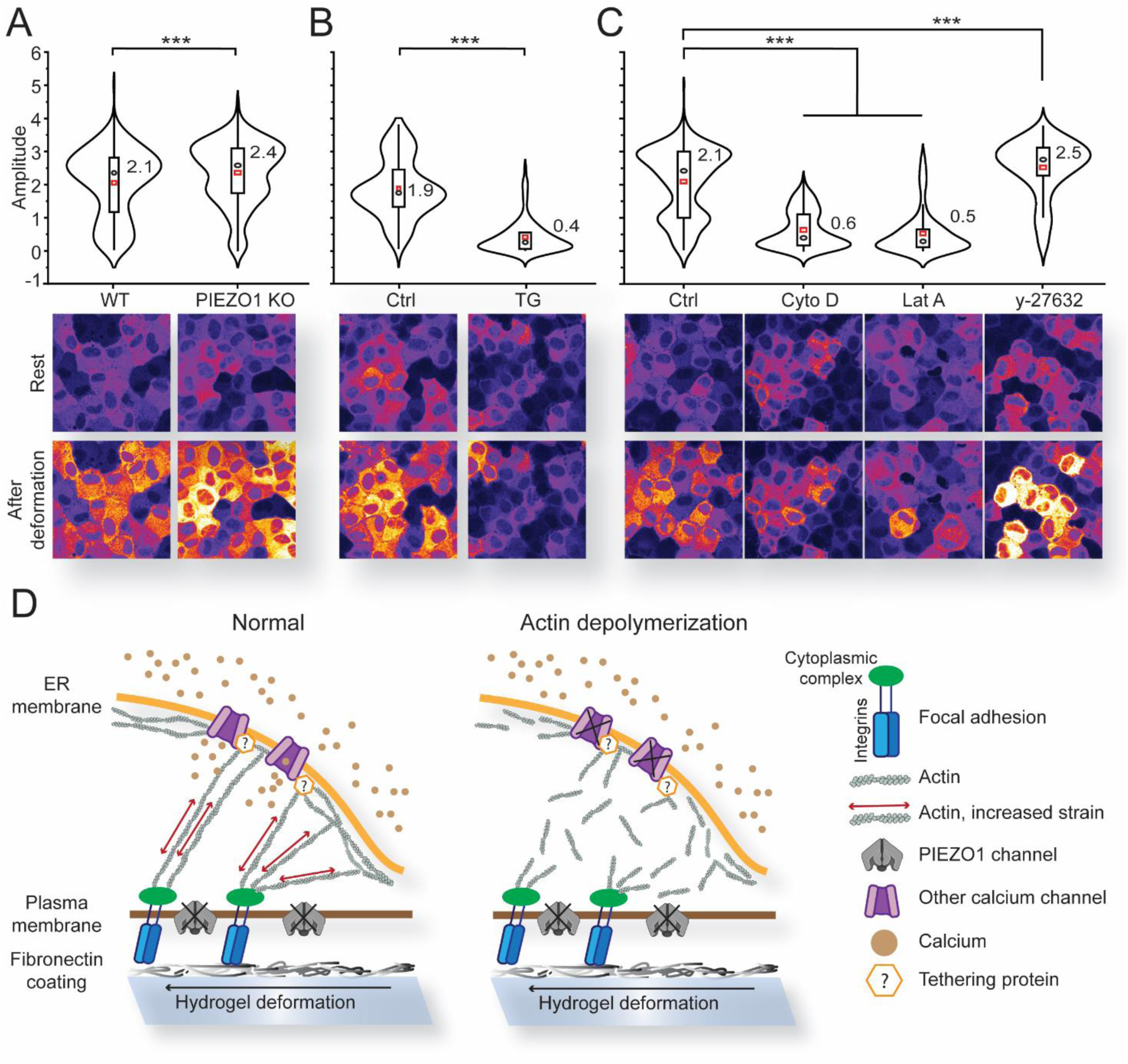
The source of calcium and roles of PIEZO1, actin, and actomyosin in mechanosensation. Above, amplitudes of deformation induced calcium signals and below, 10 slice maximum intensity projections at rest and after deformation in A) WT and PIEZO1 KO cells, B) WT cells before (ctrl) and after 15 min treatment with 1µM thapsigargin (TG), and C) before treatment (ctrl) and after 1 h treatment with 5 µg/ml cytochalasin D (Cyto D), 30 min treatment with 1 µM latrunculin A (Lat A) and 1h treatment with 10 µM y-27632. D) Schematic summary figure of the suggested mechanism: integrins of FAs adhere to the fibronectin coating on top of the hydrogel. Deformation of the hydrogel generates force to the FAs, which is transmitted to the ER via actin and possible tethering proteins. This leads to the opening of some of the calcium channels on the ER. This is disrupted if the actin cytoskeleton is depolymerized. PIEZO1 on the plasma membrane is not required in this process. In violin plots, the violin displays the density function of data points (Kernel distribution) and in the inner box, the whiskers determine the 1.5x interquartile range, the box determines the 25th and 75th percentiles and the black circles show medians and red squares the means. Mean values are noted next to the bars. *p < 0.05, **p < 0.01, and ***p < 0.001 by Students t-test (A-C).

As the signals were not triggered by the main MS ion channel on the plasma membrane, we considered that the calcium might instead be released from the ER. We used thapsigargin (TG) to inhibit the Sarco/endoplasmic reticulum Ca^2+^-ATPase (SERCA) and found that depletion of ER calcium drastically reduced cell responses to gel deformation, indicating that the ER was the main source of calcium in the detected responses (Figure 9B).

IP3Rs and STIM on the ER have been shown to require an actin tether in order to function (de Souza et al., 2021; Thillaiappan et al., 2021), and mechanical stimulation of the plasma membrane has been shown to be transmitted to the ER via actin cytoskeleton (Kim et al., 2015). We therefore examined the possible role of actomyosin machinery in the observed responses using pharmaceutical inhibitors.

The change in calcium signaling was verified to be unaffected by the solvent DMSO (Supplementary Figure 5A), and the used inhibitor concentrations and incubation times were confirmed to be effective via immunostainings of the actin cytoskeleton and phosphorylated myosin II DMSO (Supplementary Figure 5B). Once actin was depolymerized with cytochalasin D or latrunculin A, we saw significant decrease in the calcium signal amplitudes, strongly indicating that intact actin was required for the mechanical stimulation to be transmitted to the ER (Figure 9D). In contrast, inhibition myosin II with y-27632 (Rho-associated protein kinase (ROCK) inhibitor) caused no significant change, indicating that the response does not require actomyosin contractility (Figure 9C). These findings demonstrate that the effects observed with cytochalasin D and Latrunculin A are not merely due reduced cortical tension but instead suggests that actin filaments transmit forces from the cell-ECM adhesions to the ER.

## 3 Discussion

The effects of substrate viscosity and particularly elasticity as passive mechanical properties are well characterized and known to affect numerous cellular responses ranging from cell spreading to lineage specification (Chaudhuri et al., 2020; Janmey et al., 2020). However, most prior studies have focused on purely elastic materials, whereas the ECM in which cells naturally reside is viscoelastic (Chaudhuri et al., 2020). Moreover, many studies treat materials as passive entities, even though the body is a dynamic system that continuously experiences deformation as it moves and tissues reorganize. To address these two less studied phenomena, we studied calcium responses generated by ECM deformation in different viscoelastic environments. The aim of this study was to understand how the viscoelastic properties of the ECM affect cellular mechanosensitivity.

To address this question, we adapted our previously developed platform for applying basal mechanical stimulation to cells with high spatial and temporal precision (Peussa et al., 2023). By polymerizing a PAA hydrogel layer on the light responsive DR1-glass coating we could control the viscoelasticity of the cell substrate while generating basal deformation. This configuration enabled studies of how substrate viscoelasticity modulates mechanically induced calcium signaling in a MDCK monolayer.

The elasticity and viscosity of the gels were determined using AFM and rheology. The two techniques gave surprisingly different values: AFM yielded Young’s modulus values ranging from tens to hundreds of kilopascals, whereas rheology produced storage modulus values that were approximately 100-fold lower (Figure 3C−E). This is partly explained by the fact that the two techniques apply different forces (indentation vs torque) to the material and thus measure different aspects of elasticity. The techniques also apply deformation in very different ranges, as AFM functions in the nano to micrometer scale and rheology in millimeter scale. Therefore, the mechanical properties of the surface of the gel may affect AFM measurements while rheology measures the entire bulk material. For example, higher crosslinker concentration and faster polymerization reaction at the surface of the gel may result in gels with stiffer surface, which could lead to apparently higher stiffnesses values by AFM. Moreover, with AFM we were able to measure the ∼10 µm thick gels, whereas rheology measurements were restricted to macroscopic samples in the millimeter scale. While the differences between AFM and rheology measurements are large, similar differences, up to orders of magnitude, have been reported elsewhere depending on the measurement technique (Janmey et al., 2020; Wu et al., 2018). Interestingly, as the elastic modulus of a material is so strongly dependent on the type of applied force (e.g. indentation vs torque), it suggests that cells may likewise perceive materials very differently depending on the type of force they project to the ECM, thus adding another layer of complexity to understanding cell-ECM interactions.

Next, we characterized gel deformation by PIV analysis of gel-embedded fluorescent beads and by computational simulations. As expected, the ratio of deformation that reached the cell layer was inversely dependent on the gel thickness (Figure 4C−E), albeit the effect remained minimal within the thickness variation of our gels. Likewise, as in our previous work, increased laser power as well as increased number of stimulation loops resulted in larger gel deformations (Figure 4B). However, due to the large variation in deformation, no direct correlation between these parameters and gel deformation could be determined. While differences in gel thickness and heterogeneity in local elasticity (as seen in the AFM measurements) explain some variability in gel deformation, we suspect that part of the variation arises from the behavior of the DR1-glass layer. Based on our experience, DR1-glass stimulation is very sensitive to correct focusing of the stimulation laser. In our system, the focal plane for stimulation had to be determined manually, so human errors could have caused differences in DR1-glass movement. In addition, discrepancies in the thickness of the DR1-glass layer that arise from the spin coating process may have a small effect on the deformation. Lastly, as the chemistry of how PAA gel attaches to DR1-glass is unknown, it is possible that there were local differences in this attachment, which could lead to differences in how the deformation is transferred into the gel.

However, PIV measurements and simulations showed that the magnitude and dynamics of gel deformation were dependent on the gel’s viscosity and elasticity. While the stiff gels resisted DR1- glass movement more and thus underwent less deformation at the bottom of the gel, they deformed more as a solid block, thus more efficiently transmitting the deformation to the surface of the gel. Conversely, soft gels were easily displaced at the gel-DR1-glass interface, but due to their low stiffness, they transmitted less deformation to the upper surface. Moreover, in addition to the less efficient transfer of total deformation through the gel, simulations also displayed a different time scale of movement, with the stiff elastic gel deforming notably faster than the soft gels (max deformation speed 4.9 µm/s). Unsurprisingly, and in accordance with the theoretical aspects of viscosity, the deformation slowed down in soft gels as the viscosity increased (max deformation speed for soft elastic gel 2.8 µm/s and for soft viscoelastic gel 1.6 µm/s). The experimental data together with the simulations demonstrate that the viscoelastic properties of the substrate affect the magnitude and especially the dynamics of deformation. Cells may therefore receive different types of mechanical stimuli depending on their niche: lower elasticity and increased viscosity of the surrounding ECM lead to dampening of deformations and slower dynamics, and vice versa.

Finally, we studied how the viscoelastic properties of the substrate affected the mechanosensation of cells. We began by analyzing the morphology of FAs and actin stress fibers on different gel types. While the differences were smaller than anticipated, we still detected a trend in which large and elongated FAs as well as more distinguishable stress fibers were more common in cells cultured on stiff elastic gels. In accordance with previous studies, our data showed that cells developed larger and thus stronger FAs on stiff elastic gels compared to soft elastic and soft viscoelastic gels. This suggests that cells in stiff environments may also be more sensitive to mechanical stimulation due to enrichment of mechanosensitive molecules at the cell-ECM interface and increased actin anchorage. Indeed, the largest calcium signals were generated on stiff gels. Moreover, on stiff gels the mean calcium signals increased with larger deformations, suggesting that stiffness induced enrichment of FAs not only allowed mechanical cues to trigger more substantial cell responses but also enabled cells to distinguish between different force magnitudes with higher sensitivity.

Despite reports of increased viscosity making cells perceive substrates as stiffer (Chaudhuri et al., 2020; Gong et al., 2018), we did not see significant differences in FA and stress fiber morphology between the soft elastic and soft viscoelastic gels. Thus, while the substrate was in a stationary state cells did not sense the difference in viscosity of the gels used here. However, while mean calcium responses on the soft elastic gels increased slightly as deformations grew, suggesting a sensitivity to deformation magnitude (albeit less than on stiff gels), no such trend was seen on the soft viscoelastic gels. This suggests parameter beyond FA strength and deformation magnitude influence the cellular response. To understand how the detected differences in gel deformation dynamics affected the forces subjected to FAs, we expanded the computational model to simulate FA forces.

Indeed, simulations of the soft elastic and soft viscoelastic gels showed that the faster deformation of the soft elastic gels resulted in notably larger FA forces even when FA strength and deformation magnitude were kept constant. This concluded that the calcium responses were therefore dependent on the stiffness guided FA strength, the magnitude of deformation and especially the speed of deformation.

The effect of deformation dynamics is in line with our previous findings, where we showed that speed of basal shear controlled the degree of generated calcium responses (Peussa et al., 2023). In this study, where cells were grown directly on top of the DR1-glass and thus exposed to local shear, we found PIEZO1 channels to be involved in initiating these calcium responses (confirmed in Supplementary Figure 5C). We therefore suspected that the same mechanism would occur here. With this hypothesis, we studied the distribution of PIEZO1 channels, assuming to see colocalization with FAs as well as viscoelasticity dependent differences, as suggested by the literature (Jetta et al., 2023; Yao et al., 2022). TIRF imaging of MDCK II cells on glass coverslips showed strong overlap of PIEZO1 and pFAK signals, suggesting that PIEZO1 localizes at FAs on the basal membrane as expected. However, as TIRF imaging is restricted to ∼200 nm distance from the coverslip surface, this imaging technique could not be applied to gel samples. We therefore used confocal imaging. Unfortunately, due to the lower signal to noise ratio, we saw no colocalization of PIEZO1 and FAs on the gels, nor any viscoelasticity driven differences in PIEZO1 distribution or morphology. Instead, on all gel types, the channels were similarly distributed on the basal and lateral membranes. Furthermore, comparison of gel deformation induced calcium signals between WT and PIEZO1 KO cells were not biologically different, thus unexpectedly suggesting that PIEZO1 does not have a mechanistical role in the studied phenomenon.

As our current and previous data pointed out, the speed of the mechanical deformation is highly significant regarding mechanosensation and the magnitudes of generated cell responses. It is therefore possible that even slight differences in deformation time scales may activate entirely different mechanisms of mechanosensation (based on simulations, the stiff elastic gel exhibits ∼2x slower deformation than the DR1-layer). This could be further enhanced by the different deformation magnitudes (µm scale here, vs nm scale in (Peussa et al., 2023)) and the type of the deformation (displacement here vs shear in (Peussa et al., 2023)). This suggests that cells sense different magnitudes, speeds, and types of mechanical cues, resulting in distinctive cell responses. Indeed, high specificity to the type of mechanical stimulation has been widely shown. For example, electrophysiological measurements of PIEZO1 currents show distinct responses to positive and negative pressure and indentation (Coste et al., 2010; Lewis and Grandl, 2015; Pathak et al., 2014), as well as sensitivity to substrate mechanics (Bavi et al., 2019). Moreover, the type and location of mechanical stimulation has been shown generate different kinds of calcium transients in neurons (Gaub et al., 2020).

As the main MS ion channel served no mechanistic role, we suspected that the mechanosensation may be transmitted directly from FAs to the ER without need for an initial extracellular calcium influx. This was supported by treatment with thapsigargin, which showed that depleting internal calcium stores drastically decreased the calcium signals. As FAs are linked to the actin cytoskeleton which also strongly interacts with the ER, we next checked whether actin had a role in the calcium release mechanism. Multiple studies suggest that mechanical forces are transmitted from the plasma membrane to the ER in an actin and sometimes actomyosin dependent manner (Chen et al., 2025; de Souza et al., 2021; Pain et al., 2023; Thillaiappan et al., 2021). Indeed, treatment with cytochalasin D and latrunculin A, both of which lead to degradation of the actin cytoskeleton, significantly decreased the calcium signals, showing that intact actin fibers are required for the calcium release. On the contrary, inhibition of actomyosin via the ROCK activation pathway had no effect, suggesting that actomyosin contraction was not necessary. This also excludes the possibility that actin inhibitors would simply affect cortical tension and thereby make cells more pliable and less sensitive to forces. Instead, the data suggests that forces subjected to FAs are directly transmitted to the ER via actin.

Recently, multiple structures on the ER have been shown to tether to actin or to exhibit mechanosensation. IP3Rs have been shown to be linked to actin, and in fact this tether has been shown to function as an additional layer of IP3R regulation (Thillaiappan et al., 2021). While the mechanism of this regulation has not been resolved, the link to the actin cytoskeleton directly raises the possibility for mechanical sensitivity. Mechanical strain applied directly to the ER using an ontogenetic tool also revealed that MS ion channels residing at the ER open in response to this force, releasing a calcium signal (Song et al., 2024). As the ER is known to be linked to the actin cytoskeleton (Pain et al., 2023) it is possible that mechanical strain transmitted from FAs to the ER via actin could generate a similar strain on ER. While not widely studied in MDCK II cells, MS ion channels such as TRPV1 and PKD2 have been shown to release mechanically generated calcium signals at the ER (Song et al., 2024), thus making this mechanism plausible also our system. The STIM/Orai pathway has also been shown to require actin for proper function, again raising the possibility for mechanosensitive function (de Souza et al., 2021). Moreover, STIM has been identified as a tethering point between the plasma membrane and ER in addition to actin, and has been shown to transmit forces between the two membrane systems (Chen et al., 2025). We therefore suggest that the gel deformation is sensed by FAs that adhere to the fibronectin coating (Figure 10 E). These forces are transmitted directly to the ER via actin and possible tethering proteins, where the force opens a calcium channel, thus releasing a calcium signal. However, identifying the calcium channel or channels acting in this suggested pathway was not within the scope of this paper.

## 4 Conclusions

We have shown that mechanical deformation of a substrate with tunable viscoelastic properties combined to calcium imaging can hold the keys to understanding how cells sense mechanical changes in their microenvironment. The correlation between deformation speed and calcium dynamics also concurs with our previous data suggesting that the speed of basal deformation affects the degree of cell response. Moreover, our data suggests differences in time scale may even decipher which mechanosensation mechanism is activated. Our system therefore enables a new approach to mechanically stimulating cells in different viscoelastic niches for further understanding how cells integrate different kinds of mechanical stimuli.

## 5 Materials and methods

### 5.1 DR1 sample preparation

Glass coverslips (22 × 22 mm) were washed in acetone in a sonicating bath and spin coated with a solution of 3.25 % (g mL−1) of Disperse Red1 molecular glass (DR1-glass, Enamine, Ukraine) in CHCl3. The spin coating parameters were 1500 rpm for 30 s.

### 5.2 Coverslip preparation for PAA gels

22 x 22 mm coverslips spin coated with DR1-glass did not require any treatment before gel casting. However, plain bottom coverslips (used for gel thickness optimization) and top coverslips were cleaned by immersing them in 2 % Hellmanex® III (Z805939, Hellma Analytics, Müllheim, Germany) and sonicating for 30 min, followed by thorough rinsing with milliQ-H2O and ethanol. Samples were dried with a nitrogen stream. The bottom coverslips (without a DR1-glass layer) were activated with Bind-Silane (3-(Trimethoxysilyl)propyl methacrylate (TMSPMA), #M6514, Sigma-Aldrich, Saint-Louis, USA) to enable gel attachment to the coverslips. TMSPMA (0.3% (vol/vol)) and glacial acetic acid (5 % (vol/vol)) were mixed with 95 % ethanol. The solution was pipetted on coverslips and allowed to react for 3 min before rinsing the coverslips with ethanol and drying with a nitrogen stream. Top coverslips were passivated with PlusOne Repel-silane ES (Cytiva, UK) by pipetting the solution onto top coverslips and allowing to react for 5 min before rinsing with ethanol and milliQ-H2O and drying with a nitrogen stream.

### 5.3 PAA gel casting

The soft elastic (G’ = 5 kPa, G’’ = 10 Pa) and soft viscoelastic (G’ = 5 kPa, G’’ = 500 Pa) polyacrylamide gels were prepared according to (Charrier et al., 2020) except for using 1x PBS instead of H2O and the stiff elastic gels according to (Tse and Engler, 2010), using the 35 kPa gel recipe. The exact volumes are listed in Table 1.

**Table 1:** Gel recipes for making soft elastic, soft viscoelastic and stiff PAA gels with spacer beads and fluorescent beads.

| Gel | Soft elastic | Soft viscoelastic | Stiff elastic |
| --- | --- | --- | --- |
| 10x PBS (pH 7.4) | 500 µl | 500 µl | 500 µl |
| Acrylamide (40% stock) | 1000 µl | 1000 µl | 1250 µl |
| Bis (2% stock) | 250 µl | 375 µl | 750 µl |
| Linear PAA (see (Charrier et al., 2020)) | - | 2860 µl | - |
| milliQ-H<sub>2</sub>O | 3250 µl | 265 µl | 2500 µl |
| Total volume | 5000 µl | 5000 µl | 5000 µl |
| ➔ 970 µl taken to a separate tube, 20 µl fluorescent beads and 10 µl spacer beads added<br>➔ Air bubbles removed |  |  |  |
| TEMED (for 1 ml) | 2.5 µl | 2.5 µl | 2 µl |
| 10 % (W/V) APS (for 1 ml) | 7.5 µl | 7.5 µl | 10 µl |

The 5 ml mixture was mixed thoroughly. A 970 µl aliquot was taken and 20 µl of fluorescent beads (Invitrogen™ FluoSpheres™ Carboxylate-modified microspheres ⌀ 0.2 µm red (580/605) #F8810, or ⌀ 0.2 µm dark red (660/680) #F8870) and 10 µl of spacer beads (Corpuscular silicon dioxide microspheres C-SIO-5.00, #140226-10, Microspheres-Nanospheres, NY, USA) were added and carefully mixed. Air bubbles were removed with vacuum. N,N,N′,N′-Tetramethylethylenediamine (TEMED, #161-0800, Bio-Rad) and 10 % (W/V) ammonium persulphate (APS, Sigma, #A3678- 25G) were added and quickly mixed to commence polymerization. Solution was pipetted to the center of the bottom coverslip, and the top coverslip was placed on top with the Repel Silane treated surface facing the gel. The 80 g weight was carefully placed on top. We designed and 3D-printed a jig to help in placing the weight without moving the top coverslip to ensure the gels were placed in the center of the bottom coverslip (Supplementary Figure 6). Gels were allowed to polymerize in room temperature for 15 minutes. After polymerization the samples were immersed in 1 x PBS overnight, after which the top coverslips were carefully removed. Samples were stored in +4 °C in PBS.

### 5.4 Gel functionalization

Samples destined for cell culture were activated with 3,4-dihydroxy-l-phenylalanine (L-DOPA, H8502, Sigma-Aldrich) to enable functionalization with fibronectin to improve cell adhesion. L-DOPA treatment was done according to (Wouters et al., 2016). L-DOPA was dissolved in 10 mM TRIS buffer pH 10 (2 mg/ml) for 30 minutes in room temperature in dark. Once dissolved, the solution was pipetted on gels and allowed to react for 30 min in the dark, followed by two washes with PBS.

Fibronectin (purified from human plasma in house) diluted to 10 μg/ml in PBS, was pipetted on top of L-DOPA activated samples. Fibronectin was allowed to adsorb to the surface of the gel for 45 minutes. The incubation was performed under UV light to simultaneously sterilize samples.

### 5.5 Gel thickness measurements

The fluorescent beads embedded in the gels were utilized to measure gel thickness. Confocal stacks covering the entire gel volume were taken with 0.5 μm step size. Using a custom-made FIJI Macro, the confocal stacks were resliced, maximum intensity projections were taken, and Gaussian fit was performed to the fluorescence intensity profile of the projection. Two peaks in the fitted intensity profile were identified, and the gel thickness was determined as the distance between these peaks.

### 5.6 AFM and rheology

AFM measurements were performed with Bruker Icon AFM in the force volume mode using MLCT-BIO D tip (Bruker). Samples were measured in water with 50 µm x 50 µm/16 x 16 pixels scan size, using 2 µm ramp size, 1024 samples/ramp, and 1.02 Hz ramp rate. Force analysis was done using the Sneddon model on the tip approach curve using the nominal 35° tip angle and Poisson’s ratio 0.5.

Rheological characterization of the hydrogels was conducted with DHR-II hybrid rheometer (TA Instruments) equipped with 20mm parallel plate geometry. Hydrogel samples were prepared in cut 5mm syringes. First, amplitude sweeps (n=3) were conducted to determine maximum strain within linear viscoelastic region (LVE). The storage modulus (G’) of three hydrogel formulations was determined from frequency sweeps performed on three replicate samples, ranging from 0.1 Hz to 10 Hz at 22 °C with 1 % strain.

### 5.7 Gel deformation measurements

Confocal stacks of the entire gel volume were imaged before and after DR1-glass stimulation. For PIV measurements performed on plain PAA gels without cells, we used red (580/605) microspheres, as they were detectable by oculars and thus made it easier to locate the sample. In cell experiments, we used the far red (660/680) beads not to disturb the calcium indicator signal. Maximum intensity projections were taken from ∼10 slices that covered the top of the gel and bottom of the gel so that we could separately analyze deformation at both interfaces. To account for XY-directional drift, we aligned the before and after maximum intensity projections using the Stack Reg plugin in FIJI (Rigid Body). The alignment was performed based on the non-stimulated area. The PIV plugin by (Tseng et al., 2012) was used to analyze gel displacement within the stimulated ROIs.

### 5.8 Simulations of gel deformation and force generation

The simulations were conducted with the computational model presented in (Tervonen et al., 2023) with some additions and modifications. Briefly, the model depicts the apical side of epithelial cells as 2D polygons whose movements are governed by various intra- and intercellular forces. The cells are attached to an underlying hydrogel substrate, which is defined by a grid of points on its top and bottom surfaces. The apical side of the cells connects to the top surface with elastic springs that represent FAs and the vertical elasticity of the cells. Furthermore, the points on the gel top and bottom surfaces are connected by elastic springs that simulate the elasticity of the material. Finally, the bottom surface of the gel is directly attached to the underlying glass. The mechanical parameters of the cells are assumed to be independent of the gel stiffness (Rheinlaender et al., 2020), other than the spring constants describing FA, whose values were earlier fitted to gels of different stiffnesses. In addition, the cells are not entirely stationary, but include some motion as they attempt to minimize their mechanical energy. The model was tailored to describe the gel thickness used in the experiments. The movement of the cell vertices and gel points is solved using a dampened equation of motion, where the damping constants define the separate viscous behavior of the cells and the gel. The gel viscoelasticity can therefore be tuned using the damping constant and the material spring constants. To describe the displacement of the DR1-glass during stimulation with this model, the points of the bottom surface of the gel are moved within a stimulation region. The movement of the top surface of the gel and forces applied on the cells via FAs are then quantified from the simulation results. The model code is available at Github (https://github.com/atervn/epimech) and archived in Zenodo (Tervonen, 2025).

### 5.9 Cell culture

MDCK II cells expressing the genetically encoded calcium indicator jRCaMP1b (Dana et al., 2016) were maintained in Minimum Essential Medium (MEM) with GlutaMAX and Earle’s salts (41090028, Gibco), supplemented with 10 % FBS (10 500 064, Gibco). Geneticin (250 μg mL−1, G418, #4727894001, Roche, Switzerland) was used as selection antibiotic to ensure the population remained jRCaMP1b positive. Penicillin streptomycin was not used as it is a competitive inhibitor for G418. Media was changed twice a week, and cells were passaged every 7–14 days. Cells were plated at ∼3.5 × 10^4^ cells/cm^2^ density on fibronectin functionalized gel samples (see Chapter 5.4) and allowed to mature for 6 days prior to imaging. During culture on samples the media was not changed.

For calcium imaging experiments, the samples were transferred to Aireka Cells holders and conditioned media was pipetted on top.

### 5.10 DR1-glass stimulation and calcium imaging

DR1-glass stimulation and calcium imaging were performed with laser scanning confocal microscope Nikon A1R mounted in inverted Nikon Ti-E body (Nikon, Tokyo, Japan), using SR Plan Apo IR 60x water immersion objective. To enable live cell imaging, the conditions inside the microscope were set to 37 °C and 5 % CO2. A 512 x 512 pixel field of view was imaged with 200 nm pixel size. To semi-automate the imaging procedure, we utilized the JOBS module in Nikon Imaging Software (NIS) Elements. The Perfect Focus System (PFS) offset values were manually determined for the cell layer and for the DR1-glass layer (using the 561 nm autofluorescence of the DR1-glass). These values were inserted in the Stg_SetPFSLensOffsetPosition macro command to shift the focus between the DR1-glass layer and cell layer. Imaging was executed with the time lapse function and stimulation with the sequential stimulation function. Calcium imaging was performed using 561 nm excitation with 1.3 sec frame rate. Stimulation was performed with 488 nm excitation with 2.2 µsec pixel dwell time. The focus shifts and stimulation (x5) resulted in ∼13 sec blind time between the calcium imaging phases. Confocal stacks were taken before and after the calcium imaging. During cell experiments, we used dark red (660/680) microspheres as red ones would have interfered with the calcium indicator jRCamP1b.

### 5.11 Pharmaceutical testing

For pharmaceutical testing, we first performed DR1-glass stimulation experiments in control conditions in normal media. The pharmaceutical compound was added to the sample and incubated for 15 – 60 min depending on the compound: thapsigargin 1 µM 15 min (Sigma- Aldrich T9033), y- 27632 10 µM 1 h (STEMCELL Technologies 72304), ML-7 10 µM 1 h (Tocris 4310), cytochalasin D 5 µg/ml 1 h (Sigma-Aldrich C2618) and latrunculin A 1 µM 30 min (Tocris 3973). After the incubation, DR1-glass stimulation experiments were repeated. We were thus able to compare the activity of the same sample before and after treatment with the pharmaceutical substance.

### 5.12 Calcium data analysis

For each video of calcium imaging, we used Fiji ImageJ to create a 2D Z-projection image based on the standard deviation of each pixel’s intensity trace over time. Using MATLAB, we manually identified on this image the nuclei centers, around which the nuclei masks are detected by local thresholds. The signal intensity of the calcium indicator jRCaMP1b inside each nucleus throughout the time lapse video is then analyzed and extracted for features as in (Peussa et al., 2023). The 10 frames imaged before DR1-glass stimulation represented the baseline activity which was used to normalize each trace to zero. The maximum normalized signal intensity value that occurred after the first DR1-glass stimulation step was determined as the amplitude of each trace.

### 5.13 Immunostainings and image analysis

Cells were fixed with 4% paraformaldehyde (PFA, Electron Microscopy Sciences 15713-S) for 10 minutes and washed two times with PBS. Immunostainings were performed at RT protected from light. Samples were permeabilized with permeabilization buffer (0.5 % BSA and 0.5 % Triton-X 100 in PBS) for 10 min before blocking with 3 % BSA (PAN-Biotech P06-139210) in PBS for 1h. The primary antibodies (Phospho FAK pY397 [EP2160Y] ab81298, Abcam, Vinculin, clone 42H89L44, #700062, Invitrogen, and Phospho-Myosin Light Chain 2 (Ser19) Mouse mAb #3675, Cell Signaling Technology) were diluted 1:200 in blocking buffer and incubated for 1h. Samples were washed 3 x 10 min (permeabilization buffer, 1x PBS, permeabilization buffer) before adding the secondary antibody (Goat anti-Rabbit IgG (H+L) Alexa Fluor™ 488 A11008, Invitrogen, and Goat anti-mouse IgG (H+L) Alexa Fluor 488 A11001, both diluted 1:200) and phalloidin (1:100) for 1h. Samples were washed 2 x 10 min with PBS and 5 min with dH2O. DAPI (diluted 1:1000 in dH2O) was added for 3 minutes followed by washes with dH2O and PBS. Samples were either imaged without mounting by placing samples in Aireka Cells holders and immersing them in PBS or mounted with ProLong™ Diamond Antifade mountant (Invitrogen P36961). Samples were stored in +4°C protected from light.

Confocal stacks were imaged with laser scanning confocal microscope Evident FV4000 with UPLSAPO 60XS2 Silicone objective (NA 1.3). Pixel size was set to 59 x 59 nm and Z-step to 100 nm or 80 nm. Deconvolution was performed with Huygens Essential (Scientific Volume Imaging, Hilversum, Netherlands) with automated standard settings. FIJI was used to make maximum intensity projections of confocal stacks, to perform Gaussian filtering (r = 1) and to adjust image brightness and contrast.

For segmenting FAs (pFAK- and vinculin-stained samples), we took maximum intensity projections of the 3 slices where FAs were best in focus. These images were segmented and analyzed with CellProfiler™ (Stirling et al., 2021). For vinculin the diameter was set to 7–50 pixel units and for pFAK to 5−50. We used adaptive Otsu thresholding with two classes, with smoothing scale 1.3488 and correction factor 1.0. For vinculin the lower and upper bounds for thresholding were 0.05−1.0 and for pFAK 0.1−1.0. The size of adaptive window was 30. Intensity was used as a distinguishing factor for distinguishing and dividing clumped objects.

### 5.14 PIEZO1-Halo tagging, imaging and image analysis

MDCKII cells expressing PIEZO1-Halo were used for imaging PIEZO1 channels. Immunostainings were performed as described before. The Halo-tag ligand (Janelia Fluor® JFX554 HaloTag® Ligand, used at 200 nM) was incubated for 1 h along the secondaries. For TIRF imaging we used Evident-Olympus IX83 equipped with TIRF (Abbelight) and UPlanApo 100X Oil HR (NA1.50) objective (Evident). For TIRF imaging cells were cultured on fibronectin coated glass instead of gels. Confocal imaging was done with fixed and immunostained samples (as described before) by immersing unmounted samples in PBS.

### 5.15 Statistical testing

In Figure 4 the correlations between gel deformation and gel thickness, gel deformation at the top and bottom of the gel and gel deformation and standard deviation of the gel deformation were analyzed with Pearson correlation. In Figure 7 we used ANOVA for all statistical tests. In Figure 7B−D the sample sizes were so large that nonparametric tests are no longer reliable. We thus used ANOVA testing with Bonferroni post-hoc testing although the data was not normally distributed. ANOVA with Bonferroni post-hoc testing was also used for Figure 7E, as data was normally distributed. In Figure 8D−K amplitudes were not normally distributed, but we again chose to use ANOVA with Bonferroni post-hoc testing due to large sample sizes. In Figure 8L−K data was normally distributed so we used ANOVA with Bonferroni post-hoc testing.

## Acknowledgements

The authors acknowledge Tampere Imaging Facility (TIF), Biocenter Finland (BF) and Tampere University Flow Cytometry Facility (TFCF) for their service. This work made use of Tampere Microscopy Center facilities at Tampere University. Douglas Kim & GENIE Project are gratefully acknowledged for the pGP-CMV-NES-jRCaMP1b plasmid. Mari Isomäki and Samuel Peltonen are thanked for their help with sample preparation.

## References

1. Bavi, N., Richardson, J., Heu, C., Martinac, B., Poole, K., 2019. PIEZO1-Mediated Currents Are Modulated by Substrate Mechanics. ACS Nano 13, 13545–13559. 10.1021/acsnano.9b07499

2. Berridge, M.J., Lipp, P., Bootman, M.D., 2000. The versatility and universality of calcium signalling. Nat Rev Mol Cell Biol 1, 11–21. 10.1038/35036035

3. Buxboim, A., Rajagopal, K., Brown, A.E.X., Discher, D.E., 2010. How deeply cells feel: methods for thin gels. J Phys Condens Matter 22, 194116. 10.1088/0953-8984/22/19/194116

4. Chan, C.E., Odde, D.J., 2008. Traction Dynamics of Filopodia on Compliant Substrates. Science 322, 1687–1691. 10.1126/science.1163595

5. Charrier, E.E., Pogoda, K., Li, R., Park, C.Y., Fredberg, J.J., Janmey, P.A., 2020. A novel method to make viscoelastic polyacrylamide gels for cell culture and traction force microscopy. APL Bioeng 4, 036104. 10.1063/5.0002750

6. Chaudhuri, O., Cooper-White, J., Janmey, P.A., Mooney, D.J., Shenoy, V.B., 2020. Effects of extracellular matrix viscoelasticity on cellular behaviour. Nature 584, 535–546. 10.1038/s41586-020-2612-2

7. Chaudhuri, O., Gu, L., Darnell, M., Klumpers, D., Bencherif, S.A., Weaver, J.C., Huebsch, N., Mooney, D.J., 2015. Substrate stress relaxation regulates cell spreading. Nat Commun 6, 6365. 10.1038/ncomms7365

8. Chen, Z., Chen, P., Li, J., Landao-Bassonga, E., Papadimitriou, J., Gao, J., Liu, D., Tai, A., Ma, J., Lloyd, D., Kennedy, B.F., Zheng, M.H., 2025. External strain on the plasma membrane is relayed to the endoplasmic reticulum by membrane contact sites and alters cellular energetics. Sci Adv 11, eads6132. 10.1126/sciadv.ads6132

9. Coste, B., Mathur, J., Schmidt, M., Earley, T.J., Ranade, S., Petrus, M.J., Dubin, A.E., Patapoutian, A., 2010. Piezo1 and Piezo2 are essential components of distinct mechanically activated cation channels. Science 330, 55–60. 10.1126/science.1193270

10. Cox, C.D., Bavi, N., Martinac, B., 2019. Biophysical Principles of Ion-Channel-Mediated Mechanosensory Transduction. Cell Reports 29, 1–12. 10.1016/j.celrep.2019.08.075

11. Dana, H., Mohar, B., Sun, Y., Narayan, S., Gordus, A., Hasseman, J.P., Tsegaye, G., Holt, G.T., Hu, A., Walpita, D., Patel, R., Macklin, J.J., Bargmann, C.I., Ahrens, M.B., Schreiter, E.R., Jayaraman, V., Looger, L.L., Svoboda, K., Kim, D.S., 2016. Sensitive red protein calcium indicators for imaging neural activity. eLife 5, e12727. 10.7554/eLife.12727

12. de Souza, L.B., Ong, H.L., Liu, X., Ambudkar, I.S., 2021. PIP2 and septin control STIM1/Orai1 assembly by regulating cytoskeletal remodeling via a CDC42-WASP/WAVE-ARP2/3 protein complex. Cell Calcium 99, 102475. 10.1016/j.ceca.2021.102475

13. Discher, D.E., Janmey, P., Wang, Y.-L., 2005. Tissue cells feel and respond to the stiffness of their substrate. Science 310, 1139–1143. 10.1126/science.1116995

14. Doss, B.L., Pan, M., Gupta, M., Grenci, G., Mège, R.-M., Lim, C.T., Sheetz, M.P., Voituriez, R., Ladoux, B., 2020. Cell response to substrate rigidity is regulated by active and passive cytoskeletal stress. Proceedings of the National Academy of Sciences 117, 12817–12825. 10.1073/pnas.1917555117

15. Douguet, D., Honoré, E., 2019. Mammalian Mechanoelectrical Transduction: Structure and Function of Force-Gated Ion Channels. Cell 179, 340–354. 10.1016/j.cell.2019.08.049

16. Elosegui-Artola, A., Trepat, X., Roca-Cusachs, P., 2018. Control of Mechanotransduction by Molecular Clutch Dynamics. Trends in Cell Biology 28, 356–367. 10.1016/j.tcb.2018.01.008

17. Gaub, B.M., Kasuba, K.C., Mace, E., Strittmatter, T., Laskowski, P.R., Geissler, S.A., Hierlemann, A., Fussenegger, M., Roska, B., Müller, D.J., 2020. Neurons differentiate magnitude and location of mechanical stimuli. Proceedings of the National Academy of Sciences 117, 848– 856. 10.1073/pnas.1909933117

18. Gong, Z., Szczesny, S.E., Caliari, S.R., Charrier, E.E., Chaudhuri, O., Cao, X., Lin, Y., Mauck, R.L., Janmey, P.A., Burdick, J.A., Shenoy, V.B., 2018. Matching material and cellular timescales maximizes cell spreading on viscoelastic substrates. Proceedings of the National Academy of Sciences 115, E2686–E2695. 10.1073/pnas.1716620115

19. Iskratsch, T., Wolfenson, H., Sheetz, M.P., 2014. Appreciating force and shape — the rise of mechanotransduction in cell biology. Nat Rev Mol Cell Biol 15, 825–833. 10.1038/nrm3903

20. Janmey, P.A., Fletcher, D.A., Reinhart-King, C.A., 2020. Stiffness Sensing by Cells. Physiol Rev 100, 695–724. 10.1152/physrev.00013.2019

21. Jetta, D., Shireen, T., Hua, S.Z., 2023. Epithelial cells sense local stiffness via Piezo1 mediated cytoskeletal reorganization. Front Cell Dev Biol 11, 1198109. 10.3389/fcell.2023.1198109

22. Kim, T.-J., Joo, C., Seong, J., Vafabakhsh, R., Botvinick, E.L., Berns, M.W., Palmer, A.E., Wang, N., Ha, T., Jakobsson, E., Sun, J., Wang, Y., 2015. Distinct mechanisms regulating mechanical force-induced Ca2+ signals at the plasma membrane and the ER in human MSCs. eLife 4, e04876. 10.7554/eLife.04876

23. Kirby, R., Georges Sabat, R., Nunzi, J.-M., Lebel, O., 2014. Disperse and disordered: a mexylaminotriazine-substituted azobenzene derivative with superior glass and surface relief grating formation. Journal of Materials Chemistry C 2, 841–847. 10.1039/C3TC32034K

24. Lewis, A.H., Grandl, J., 2015. Mechanical sensitivity of Piezo1 ion channels can be tuned by cellular membrane tension. eLife 4, e12088. 10.7554/eLife.12088

25. Mai, Z., Lin, Y., Lin, P., Zhao, X., Cui, L., 2024. Modulating extracellular matrix stiffness: a strategic approach to boost cancer immunotherapy. Cell Death Dis 15, 307. 10.1038/s41419-024-06697-4

26. Martino, F., Perestrelo, A.R., Vinarský, V., Pagliari, S., Forte, G., 2018. Cellular Mechanotransduction: From Tension to Function. Front. Physiol. 9. 10.3389/fphys.2018.00824

27. Pain, C., Tolmie, F., Wojcik, S., Wang, P., Kriechbaumer, V., 2023. intER-ACTINg: The structure and dynamics of ER and actin are interlinked. Journal of Microscopy 291, 105–118. 10.1111/jmi.13139

28. Pathak, M.M., Nourse, J.L., Tran, T., Hwe, J., Arulmoli, J., Le, D.T.T., Bernardis, E., Flanagan, L.A., Tombola, F., 2014. Stretch-activated ion channel Piezo1 directs lineage choice in human neural stem cells. Proc Natl Acad Sci U S A 111, 16148–16153. 10.1073/pnas.1409802111

29. Peussa, H., Fedele, C., Tran, H., Marttinen, M., Fadjukov, J., Mäntylä, E., Priimägi, A., Nymark, S., Ihalainen, T.O., 2023. Light-Induced Nanoscale Deformation in Azobenzene Thin Film Triggers Rapid Intracellular Ca2+ Increase via Mechanosensitive Cation Channels. Advanced Science 10, 2206190. 10.1002/advs.202206190

30. Plotnikov, S.V., Pasapera, A.M., Sabass, B., Waterman, C.M., 2012. Force Fluctuations within Focal Adhesions Mediate ECM-Rigidity Sensing to Guide Directed Cell Migration. Cell 151, 1513– 1527. 10.1016/j.cell.2012.11.034

31. Plotnikov, S.V., Sabass, B., Schwarz, U.S., Waterman, C.M., 2014. High-Resolution Traction Force Microscopy. Methods Cell Biol 123, 367–394. 10.1016/B978-0-12-420138-5.00020-3

32. Poole, K., Herget, R., Lapatsina, L., Ngo, H.-D., Lewin, G.R., 2014. Tuning Piezo ion channels to detect molecular-scale movements relevant for fine touch. Nat Commun 5, 3520. 10.1038/ncomms4520

33. Ranade, S.S., Syeda, R., Patapoutian, A., 2015. Mechanically Activated Ion Channels. Neuron 87, 1162–1179. 10.1016/j.neuron.2015.08.032

34. Rheinlaender, J., Dimitracopoulos, A., Wallmeyer, B., Kronenberg, N.M., Chalut, K.J., Gather, M.C., Betz, T., Charras, G., Franze, K., 2020. Cortical cell stiffness is independent of substrate mechanics. Nat. Mater. 19, 1019–1025. 10.1038/s41563-020-0684-x

35. Rocio Servin-Vences, M., Moroni, M., Lewin, G.R., Poole, K., 2017. Direct measurement of TRPV4 and PIEZO1 activity reveals multiple mechanotransduction pathways in chondrocytes. eLife 6, e21074. 10.7554/eLife.21074

36. Song, Y., Zhao, Z., Xu, L., Huang, P., Gao, J., Li, J., Wang, X., Zhou, Y., Wang, J., Zhao, W., Wang, L., Zheng, C., Gao, B., Jiang, L., Liu, K., Guo, Y., Yao, X., Duan, L., 2024. Using an ER-specific optogenetic mechanostimulator to understand the mechanosensitivity of the endoplasmic reticulum. Developmental Cell 59, 1396–1409.e5. 10.1016/j.devcel.2024.03.014

37. Stirling, D.R., Swain-Bowden, M.J., Lucas, A.M., Carpenter, A.E., Cimini, B.A., Goodman, A., 2021. CellProfiler 4: improvements in speed, utility and usability. BMC Bioinformatics 22, 433. 10.1186/s12859-021-04344-9

38. Tervonen, A., 2025. atervn/epimech: v1.1.1. 10.5281/zenodo.16892400

39. Tervonen, A., Korpela, S., Nymark, S., Hyttinen, J., Ihalainen, T.O., 2023. The Effect of Substrate Stiffness on Elastic Force Transmission in the Epithelial Monolayers over Short Timescales. Cel. Mol. Bioeng. 16, 475–495. 10.1007/s12195-023-00772-0

40. Thillaiappan, N.B., Smith, H.A., Atakpa-Adaji, P., Taylor, C.W., 2021. KRAP tethers IP3 receptors to actin and licenses them to evoke cytosolic Ca2+ signals. Nat Commun 12, 4514. 10.1038/s41467-021-24739-9

41. Tse, J.R., Engler, A.J., 2010. Preparation of Hydrogel Substrates with Tunable Mechanical Properties. Current Protocols in Cell Biology 47, 10.16.1-10.16.16. 10.1002/0471143030.cb1016s47

42. Tseng, Q., Duchemin-Pelletier, E., Deshiere, A., Balland, M., Guillou, H., Filhol, O., Théry, M., 2012. Spatial organization of the extracellular matrix regulates cell–cell junction positioning. Proceedings of the National Academy of Sciences 109, 1506–1511. 10.1073/pnas.1106377109

43. Tumenbayar, B.-I., Pham, K., Biber, J.C., Tutino, V.M., Brazzo III, J.A., Yao, P., Bae, Y., 2025. FAK and p130Cas Modulate Stiffness-Mediated Early Transcription and Cellular Metabolism. Cytoskeleton 82, 197–215. 10.1002/cm.21971

44. Wilson, D.I., 2018. What is rheology? Eye (Lond) 32, 179–183. 10.1038/eye.2017.267

45. Wouters, O.Y., Ploeger, D.T.A., van Putten, S.M., Bank, R.A., 2016. 3,4-Dihydroxy-L-Phenylalanine as a Novel Covalent Linker of Extracellular Matrix Proteins to Polyacrylamide Hydrogels with a Tunable Stiffness. Tissue Engineering. Part C: Methods 22, 91–101. 10.1089/ten.tec.2015.0312

46. Wu, P.-H., Aroush, D.R.-B., Asnacios, A., Chen, W.-C., Dokukin, M.E., Doss, B.L., Durand-Smet, P., Ekpenyong, A., Guck, J., Guz, N.V., Janmey, P.A., Lee, J.S.H., Moore, N.M., Ott, A., Poh, Y.-C., Ros, R., Sander, M., Sokolov, I., Staunton, J.R., Wang, N., Whyte, G., Wirtz, D., 2018. A comparison of methods to assess cell mechanical properties. Nat Methods 15, 491–498. 10.1038/s41592-018-0015-1

47. Yao, M., Tijore, A., Cheng, D., Li, J.V., Hariharan, A., Martinac, B., Tran Van Nhieu, G., Cox, C.D., Sheetz, M., 2022. Force- and cell state–dependent recruitment of Piezo1 drives focal adhesion dynamics and calcium entry. Science Advances 8, eabo1461. 10.1126/sciadv.abo1461

48. Yeh, Y.-C., Ling, J.-Y., Chen, W.-C., Lin, H.-H., Tang, M.-J., 2017a. Mechanotransduction of matrix stiffness in regulation of focal adhesion size and number: reciprocal regulation of caveolin-1 and β1 integrin. Sci Rep 7, 15008. 10.1038/s41598-017-14932-6

49. Yeh, Y.-C., Ling, J.-Y., Chen, W.-C., Lin, H.-H., Tang, M.-J., 2017b. Mechanotransduction of matrix stiffness in regulation of focal adhesion size and number: reciprocal regulation of caveolin-1 and β1 integrin. Sci Rep 7, 15008. 10.1038/s41598-017-14932-6

50. Zhou, D.W., Lee, T.T., Weng, S., Fu, J., García, A.J., 2017. Effects of substrate stiffness and actomyosin contractility on coupling between force transmission and vinculin–paxillin recruitment at single focal adhesions. MBoC 28, 1901–1911. 10.1091/mbc.e17-02-0116

51. Zimerman, B., Volberg, T., Geiger, B., 2004. Early molecular events in the assembly of the focal adhesion-stress fiber complex during fibroblast spreading. Cell Motility 58, 143–159. 10.1002/cm.20005

