## Supplementary figures for "Viscoelasticity driven deformation dynamics of the substrate affect mechanically induced Ca^2+^ signals"

### Supplementary Figures for:
### Viscoelasticity driven deformation dynamics of the substrate affect mechanically induced $Ca^{2+}$ signals

Peussa H, Peltola S*, Tervonen A*, Lehtimäki S, Kauppila M, Bhati R, Fedele C, Tran H, Mäntylä E, Priimägi A, Nymark S, Ihalainen TO.

* equal contribution

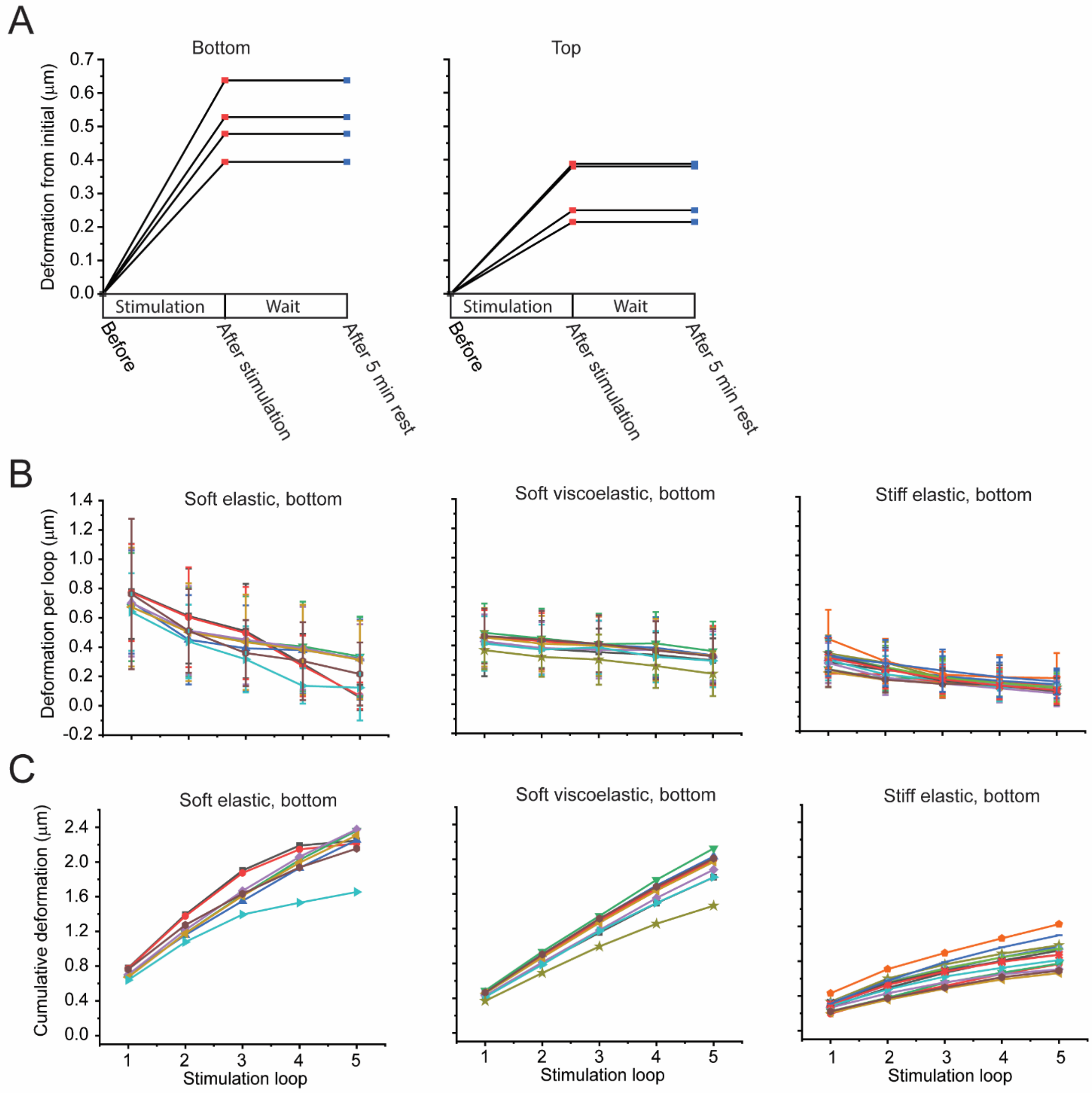


***Supplementary Figure 1: Stability and cumulation of deformation.*** *A) Gel deformation after stimulation and after five minutes rest on the bottom and top of the gel. B) Mean ± standard deviation of deformation at the bottom of the gel after each stimulation loop for soft elastic, soft viscoelastic and stiff elastic gels. C) Cumulative deformation from loop to loop for each gel type.*

***Supplementary Figure 2***: *Montages of pFAK and vinculin on soft elastic, soft viscoelastic and stiff elastic gels. Montages present maximum intensity projections over the basal side of cells, and the segmented versions of the same images. Scale bars 10 µm.*

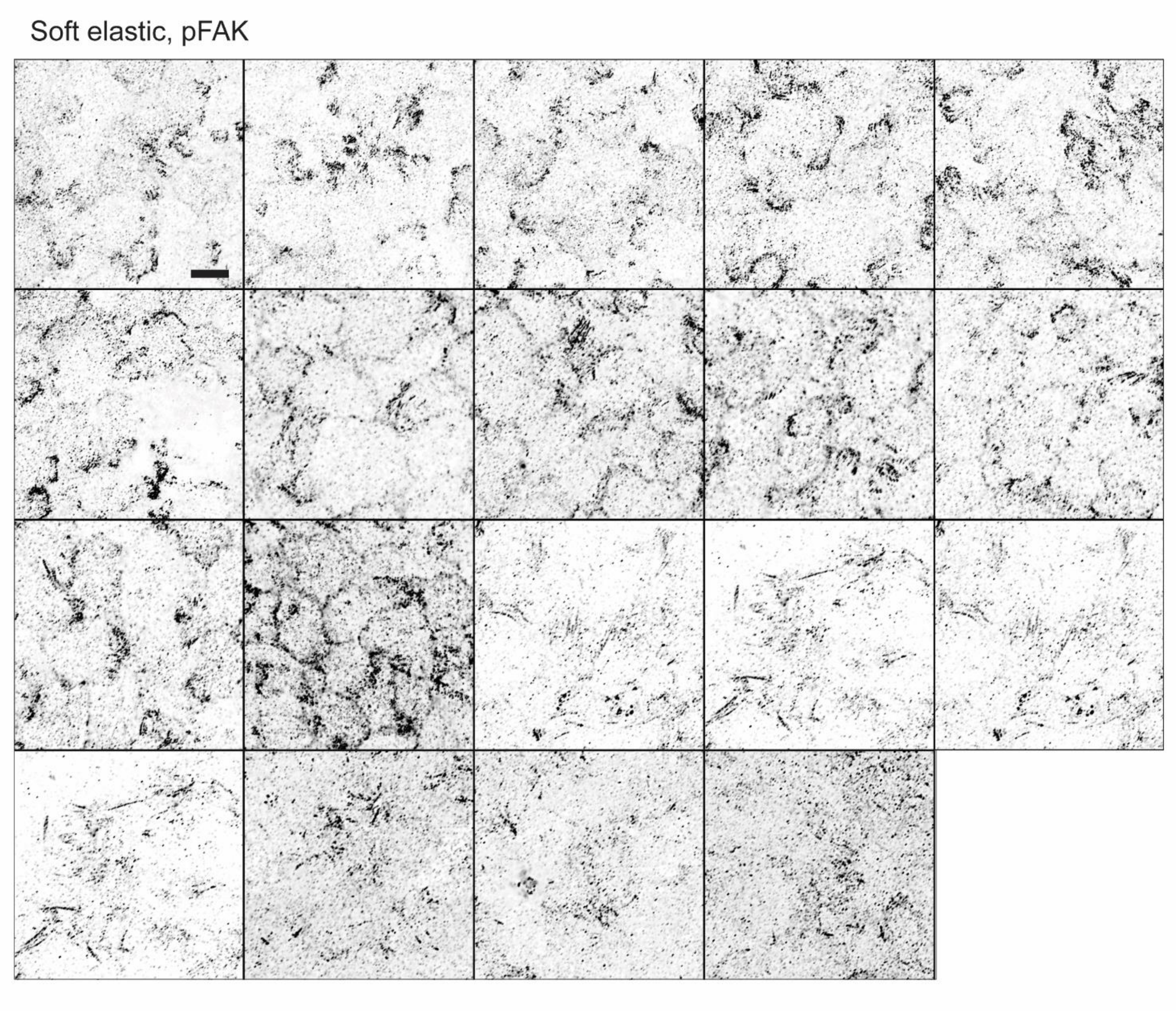

Soft elastic, pFAK, segmented

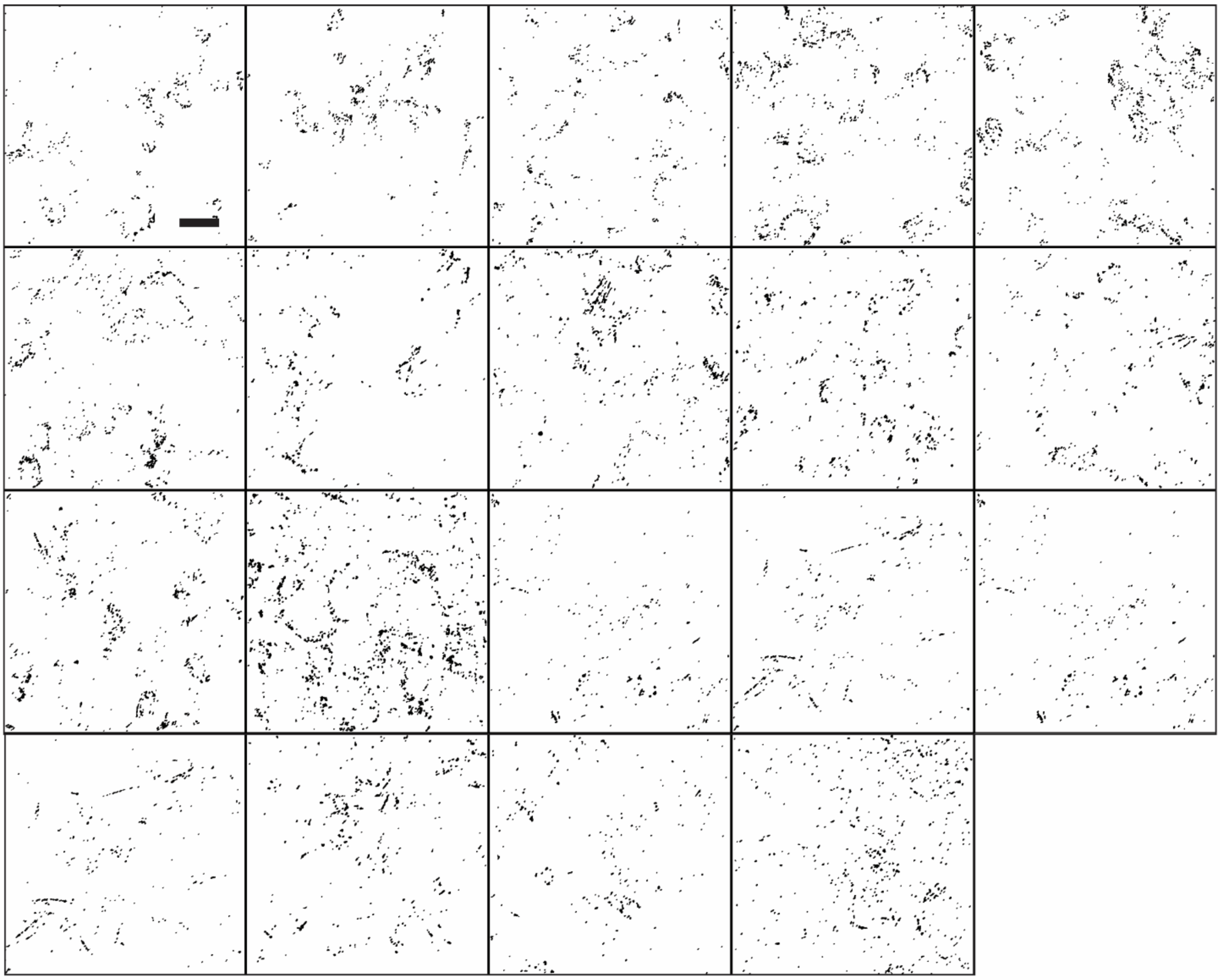

Soft viscoelastic, pFAK

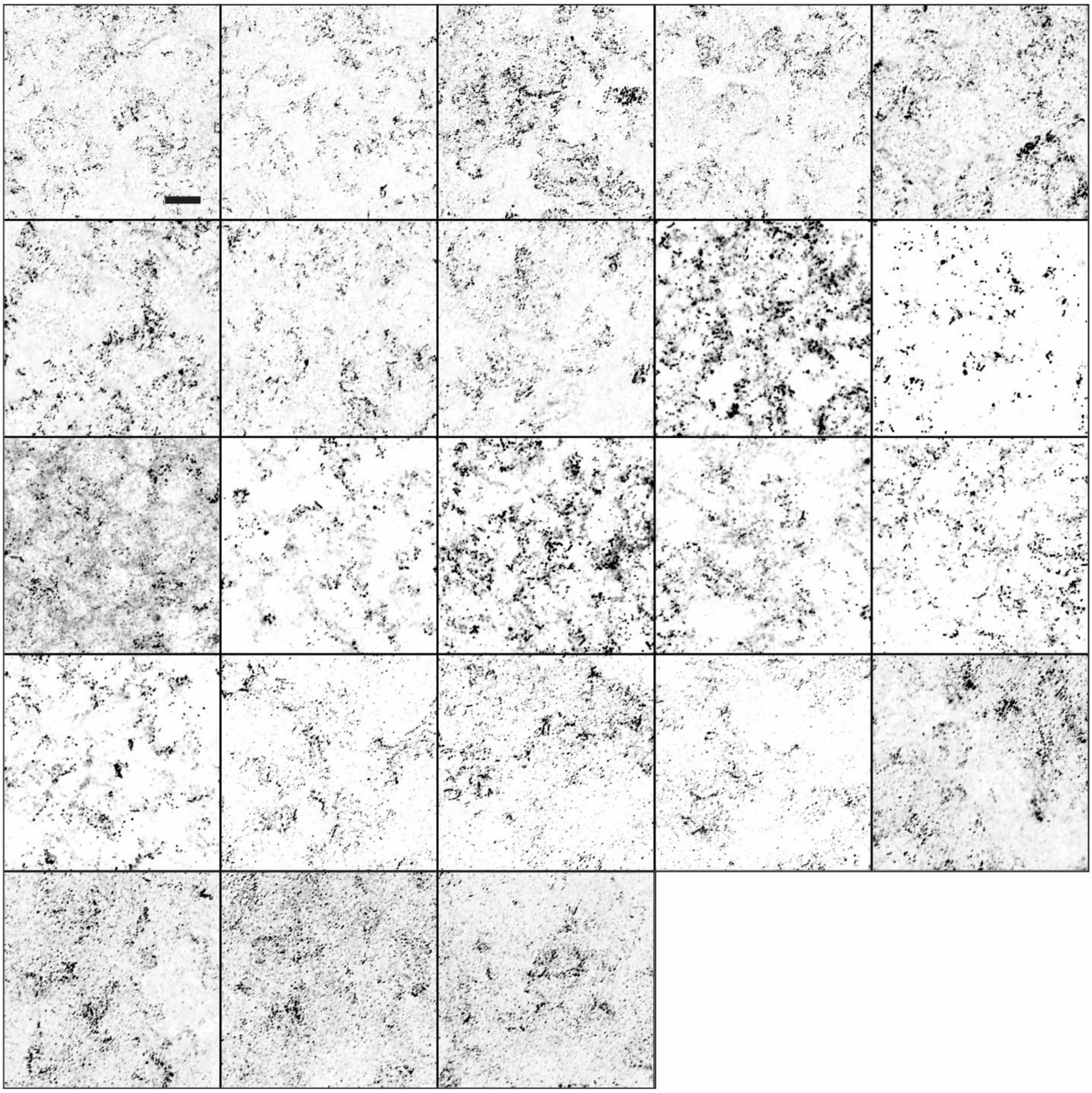

Soft viscoelastic, pFAK, segmented

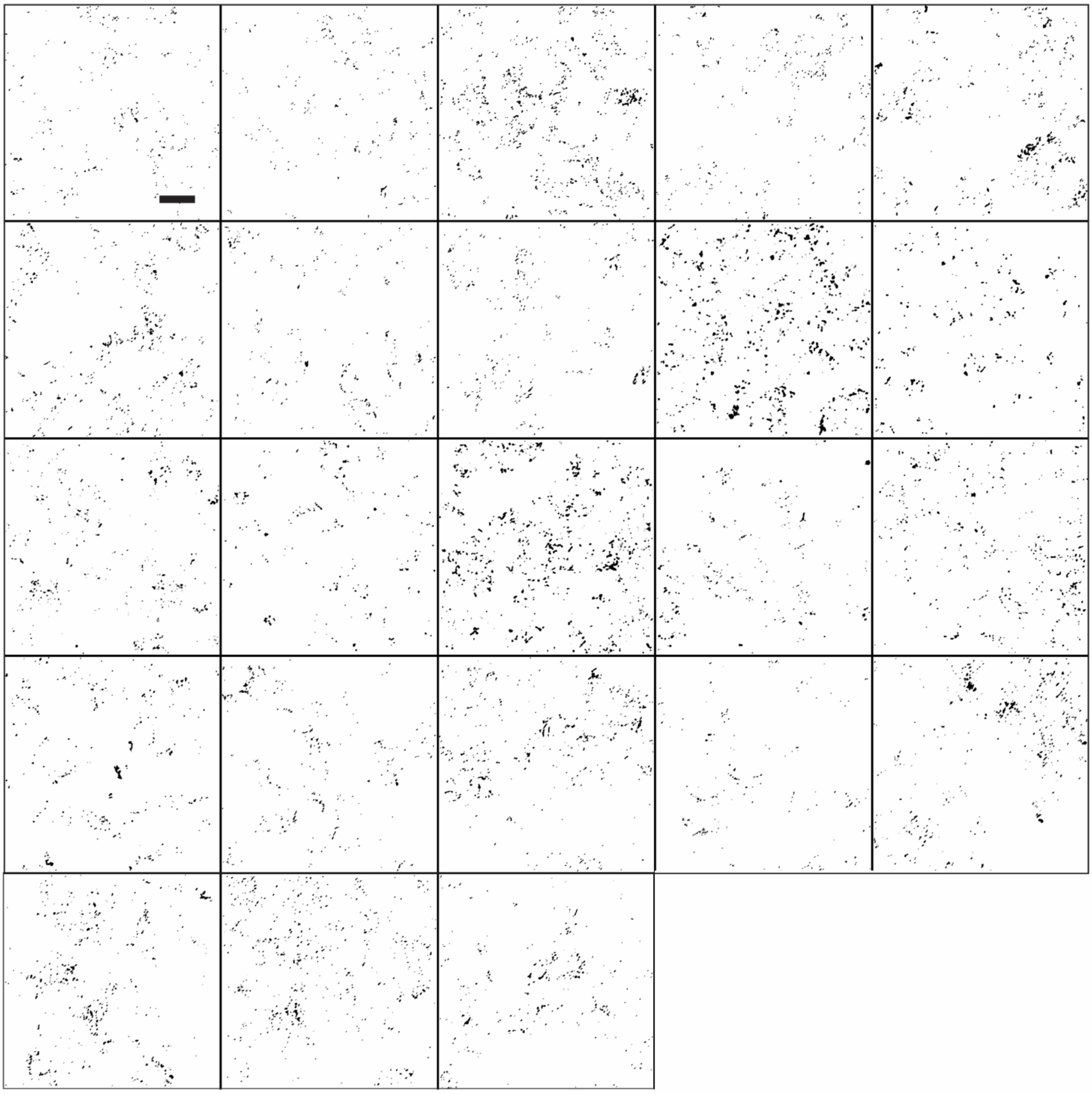

Stiff elastic, pFAK

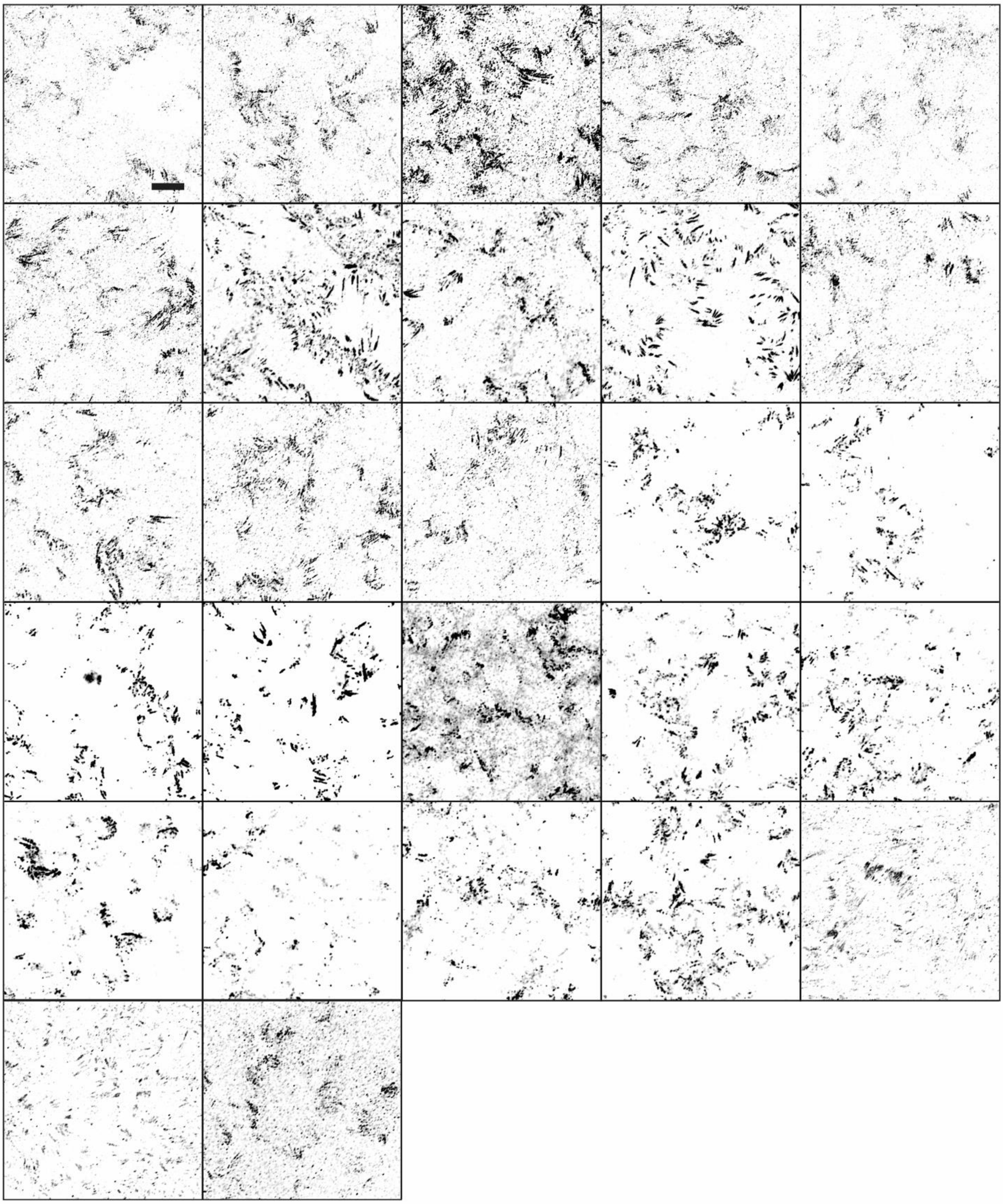

Stiff elastic, pFAK, segmented

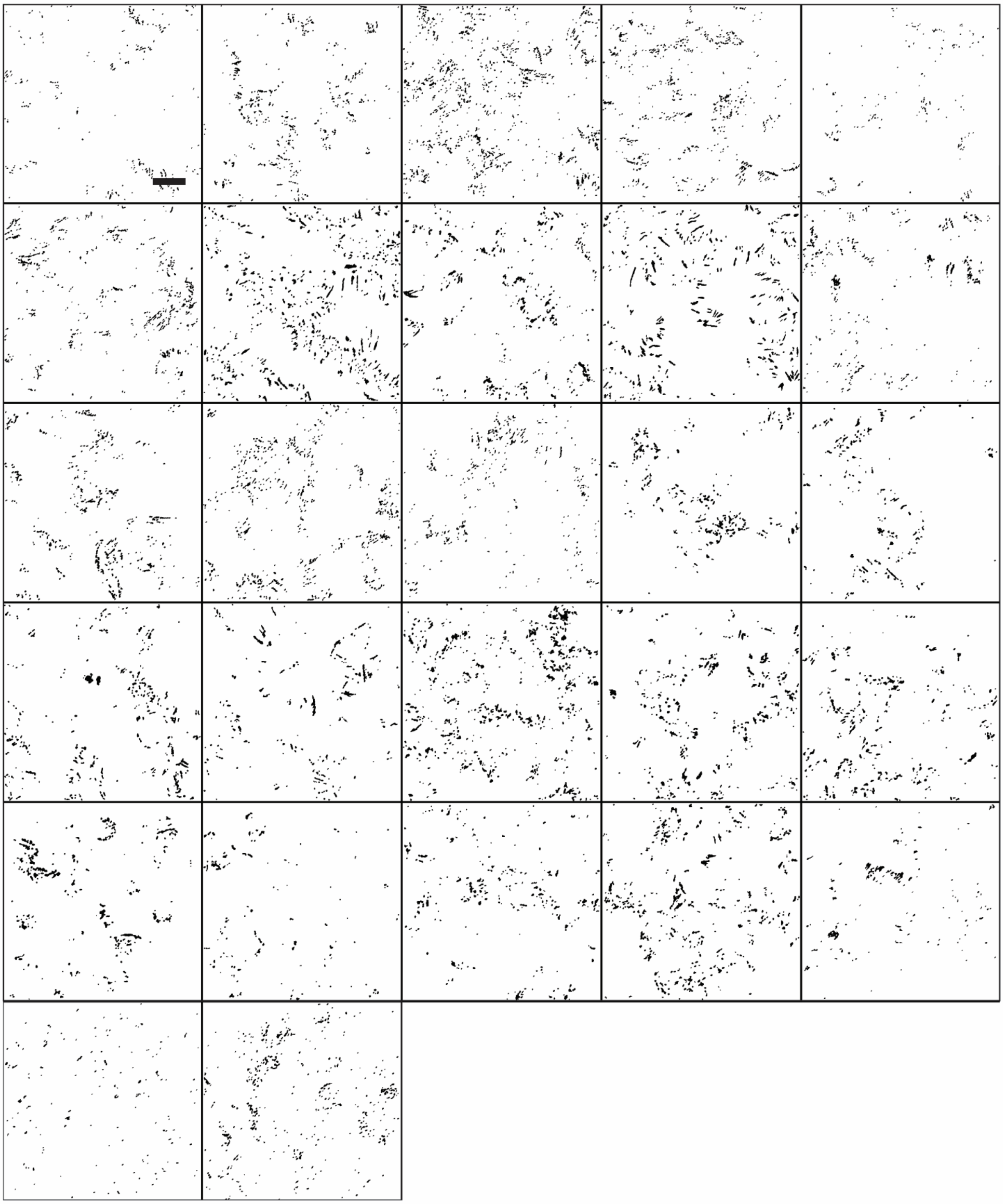

Soft elastic, vinculin

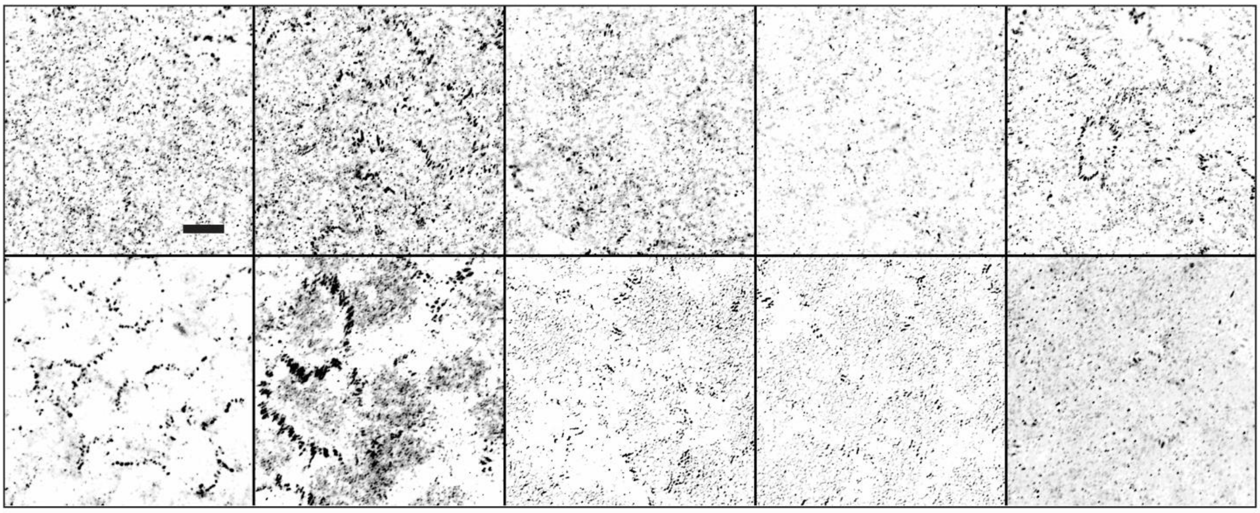

Soft elastic, vinculin, segmented

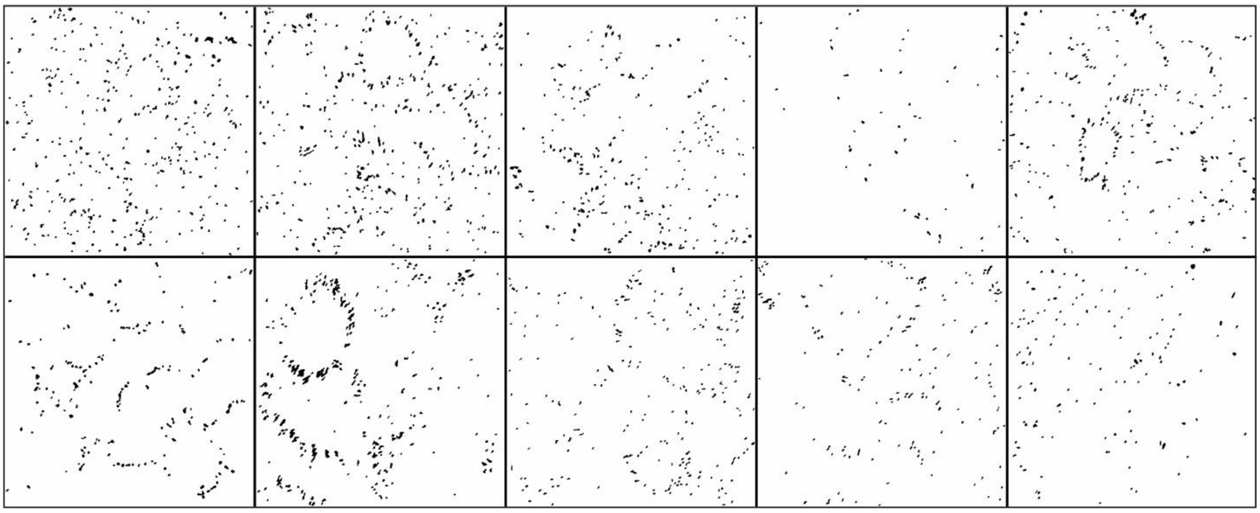

Soft viscoelastic, vinculin

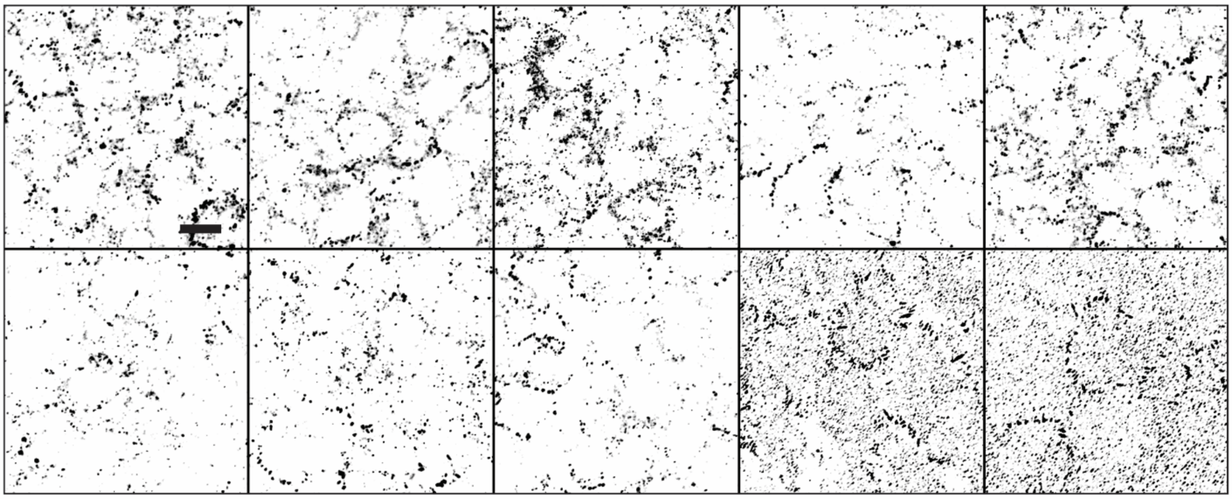

Soft viscoelastic, vinculin, segmented

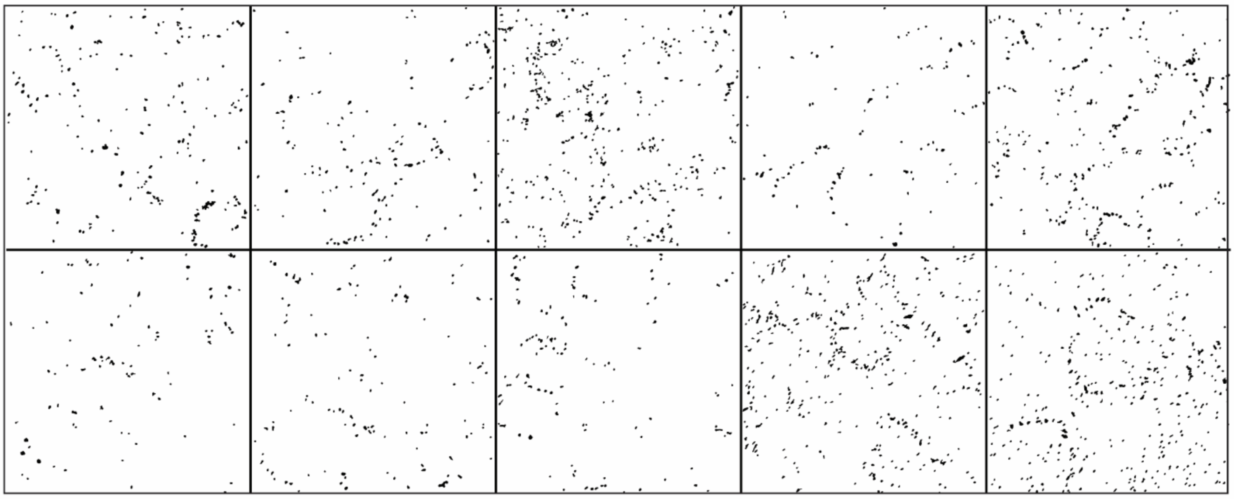

Stiff elastic, vinculin

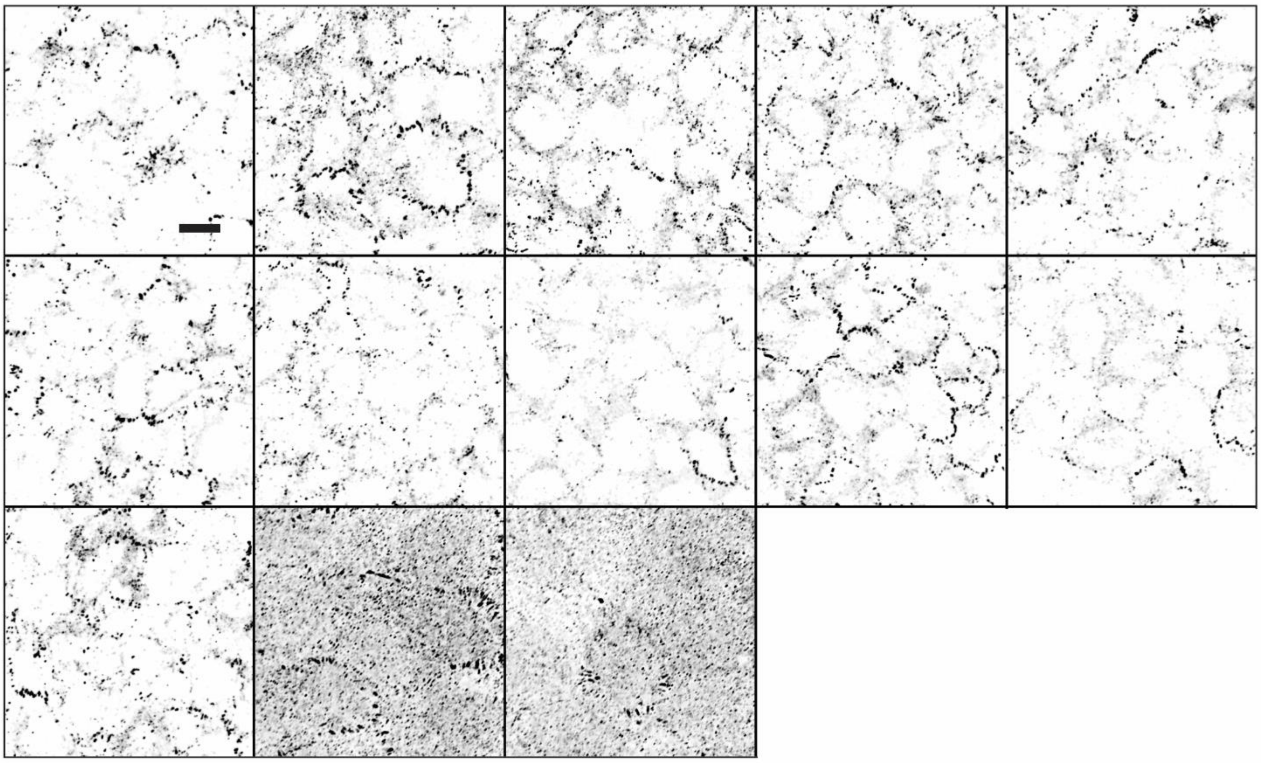

Stiff elastic, vinculin, segmented

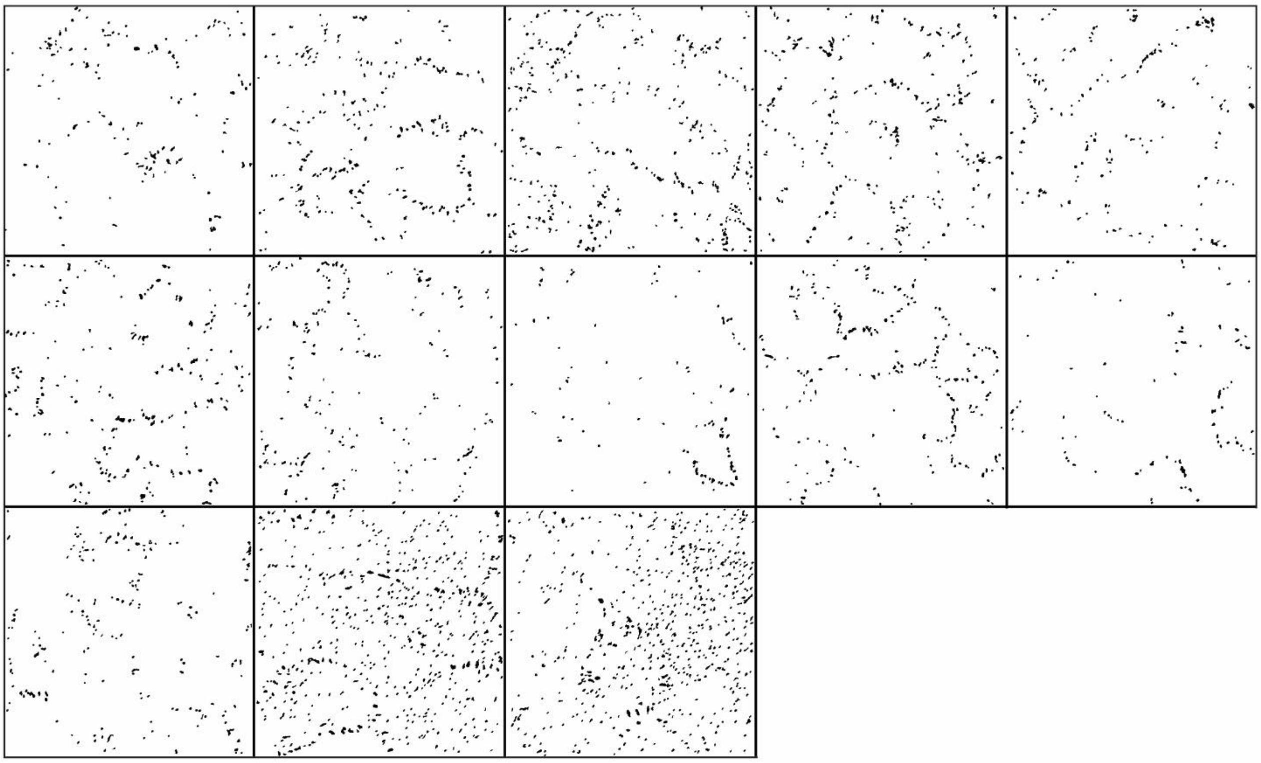

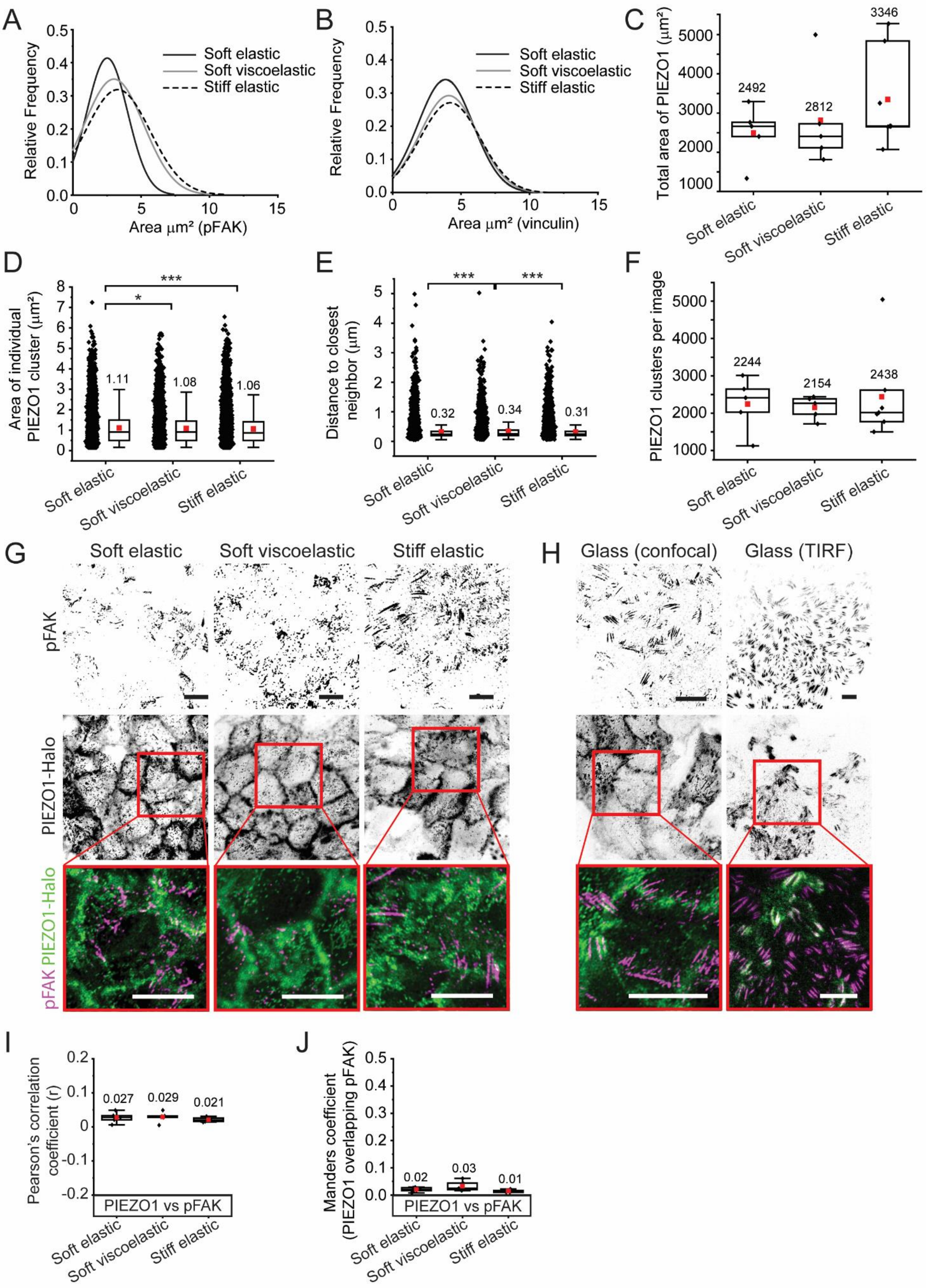


***Supplementary Figure 3: Focal adhesion area distribution, PIEZO1 channel expression and colocalization with pFAK.*** *A) Distribution of FA areas according to pFAK immunostainings and B) vinculin immunostainings. C) Total area of PIEZO1 signal per image. D) Area of individual PIEZO1 clusters, E) Shortest distance between PIEZO1 clusters. F) Number of PIEZO1 clusters per image. G) Basal PIEZO1 and pFAK signal on the different gel types. Scale bars 10 µm. H) Comparison of basal PIEZO1 and pFAK signal imaged by TIRF or confocal microscopy. Scale bars 10 µm. I) Pearson's correlation coefficient of signal intensities of PIEZO1 and pFAK and J) Manders coefficient of PIEZO1 signal overlapping with pFAK signal. In box plots, the whiskers determine the 5th and 95th percentiles, and the box determines the 25th and 75th percentiles. Means are marked with red squares, with mean values noted next to the bars. The vertical line represents the median value. *p < 0.05, **p < 0.01, and ***p < 0.001 by one-way ANOVA with Bonferroni Post Hoc test (D,E).*

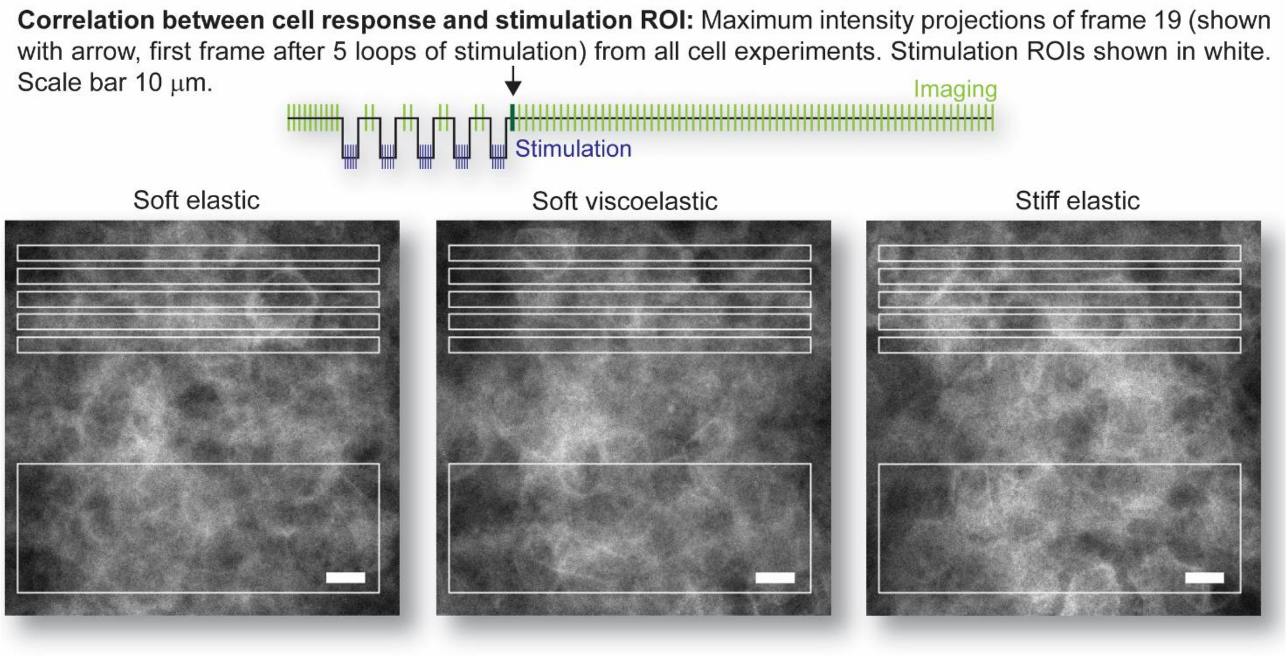


***Supplementary Figure 4: Correlation between cell response and stimulation ROI.*** *Maximum intensity projections of frame 19 (shown with arrow, first frame after 5 loops of stimulation) from all cell experiments. Stimulation ROIs shown in white. Scale bar 10 µm. Signal intensity does not correlate with stimulation ROIs.*

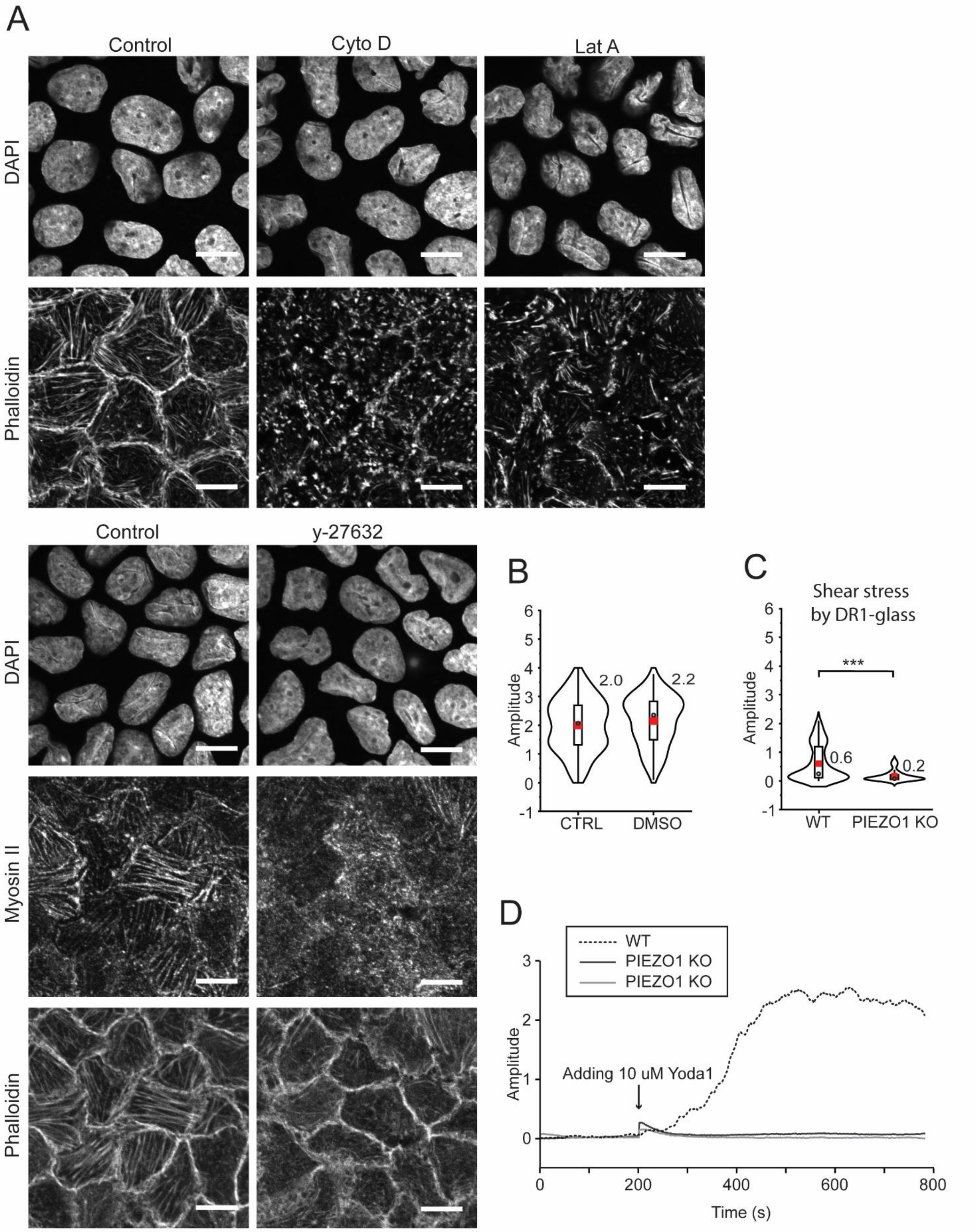


***Supplementary Figure 5: Controls for pharmaceutical experiments.*** *A) 10 slice maximum intensity projections of the basal side of cells showing the nuclei and phalloidin in control cells (10 µM DMSO) and in cells after treatment with Cyto D or Lat A, and nuclei, phosphorylated myosin II and phalloidin in control cells and in cells after treatment with y-27632. Scale bars 10 µm. B) Amplitudes of calcium signals after mechanical stimulation in normal media (CTRL) and in media containing 10 µM DMSO. C) Amplitudes of calcium signals in WT and PIEZO1 KO cells after mechanical stimulation by direct shear from DR1-glass. D) Recorded calcium signals from two PIEZO1 KO samples (black and grey solid lines) and one WT sample (black dashed line) showing the response to 10 µM Yoda1. Traces are normalized to zero with the baseline activity recorded before adding Yoda1. In violin plots, the violin displays the density function of data points (Kernel distribution) and in the inner box, the whiskers determine the 1.5x interquartile range, the box determines the 25th and 75th percentiles and the black circles show medians and red squares the means. Mean values are noted next to the bars. *p < 0.05, **p < 0.01, and ***p < 0.001 by Students t-test (B-C).*

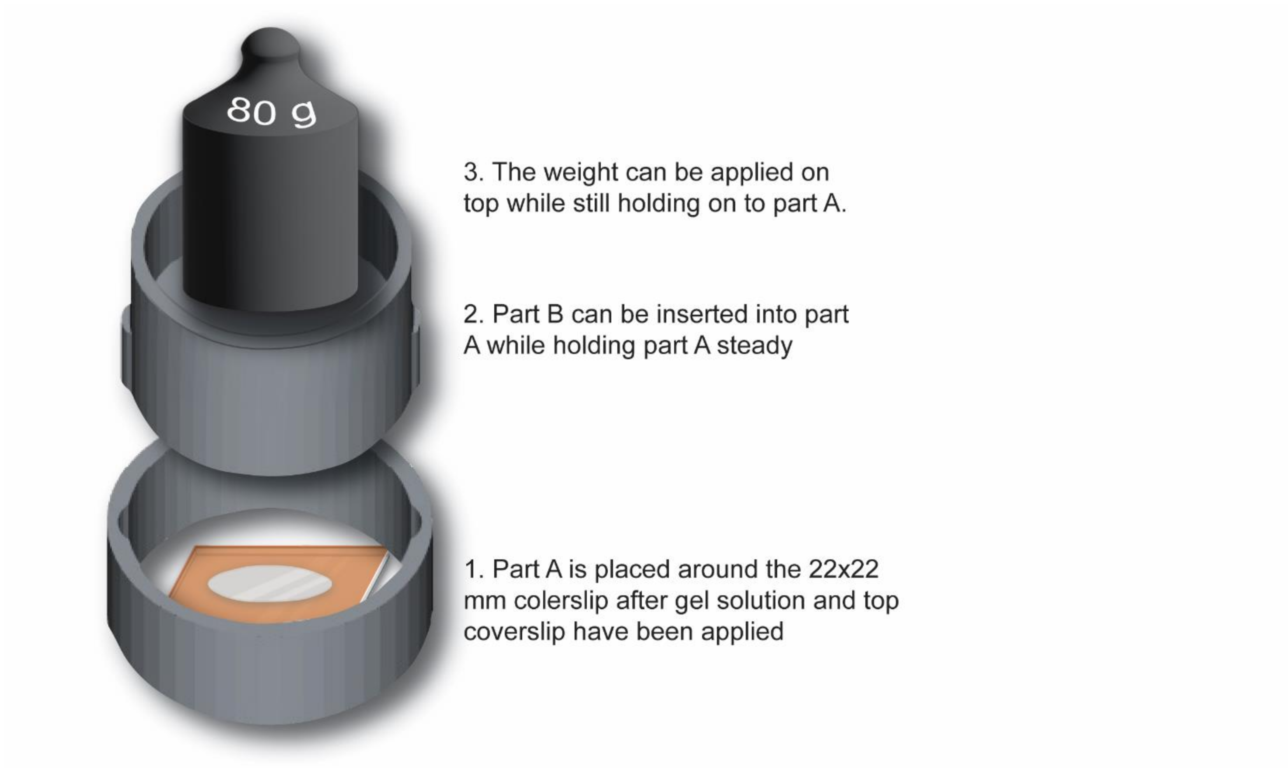


***Supplementary Figure 6****: 3D-printed jig to help with gel casting.*
